# Ketone bodies mitigate against systemic inflammation-induced changes in brain energy metabolism and delirium-like deficits in aged mice

**DOI:** 10.64898/2026.01.22.701114

**Authors:** Pierre-Louis Hollier, M.K. Kirsten Chui, Javier Cuitavi, Paul Denver, Hugh J Delaney, John C. Newman, Colm Cunningham

**Affiliations:** School of Biochemistry and Immunology, Trinity Biomedical Sciences Institute; Trinity College Institute of Neuroscience, Trinity College Dublin, Rep. Of Ireland; Buck Institute for Research on Aging, Novato, CA, United States; Division of Geriatrics, University of California San Francisco, San Francisco, CA, United States

**Keywords:** Systemic inflammation, delirium, ketone, ketogenesis, ageing, aging, beta hydroxybutyrate, sickness behaviour, hypoglycemia, Energy metabolism, ketone body, BHB, delirium, aging, brain, LPS

## Abstract

Acute systemic inflammation affects brain function, with detrimental consequences in aged individuals. These include delirium, an acute neuropsychiatric syndrome characterized by fluctuating disturbances in attention, perception and cognition. Delirium is associated with disrupted brain energy metabolism but our understanding of this during acute systemic inflammation is limited. Here we hypothesized that LPS-induced systemic inflammation would disrupt brain energy metabolism in aged C57BL6J mice and that the consequent functional impairments would be mitigated by ketone body utilization. We investigated ketone body effects in sickness behaviour, inflammation, energy metabolism and cognitive function. Real-time changes in utilisation of energy sources were quantified by indirect calorimetry and administration of radioisotope-labelled glucose and betahydroxybutyrate. Mass-spectrometry metabolomics was used to index severity of behavioural distrurbances to changes in hippocampal energy metabolism. LPS precipitated hypoglycemia and induced a whole-body switch from carbohydrate to lipid utilisation. Despite this, hippocampal insulin resistance and preserved brain glucose was observed while alternative carbohydrates, mannose and fructose, became depleted. Ketone ester treatment reversed insulin resistance, mitigated sickness behaviour and prevented delirium-like cognitive dysfunction without altering pro-inflammatory responses. Our results show that promoting ketone body usage mitigates systemic inflammation-induced brain energy disruption and prevents delirium-like cognitive deficits in aged mice.

## Introduction

An episode of viral or bacterial infection triggers a variety of behavioral, metabolic and neuroendocrine changes, encompassing lethargy, hypophagia, hypo– or hyperthermia, anhedonia and withdrawal from social interaction. Such changes are part of a phenomenon called sickness behaviour, an adaptive strategy to prioritise fighting the infection while preserving essential functions of the organism. Systemic administration of lipopolysaccharide (LPS), a component of the outer membrane of gram-negative bacteria, has been widely used to induce sickness behaviour in rodents^1^ and humans^2^. Several other agonists of toll-like recpetors and indeed pro-inflammatory cytokines such as IL-1β and TNFα can also induce similar responses^3–5^. These responses are regarded as adaptive in the process of restoring homeostasis following infection, but it is known that acute systemic inflammation induces exaggerated sickness behaviour responses when occurring in older individuals^6^ and, clinically, can trigger deleterious consequences such as delirium^7^.

Delirium is a complex neuropsychiatric syndrome characterized by an acute onset of fluctuating disturbances in attention, awareness, perception, and cognition^8^ that happens more frequently in older adults and in patients with pre-existing neuropathology^9,10^. Our understanding of the pathophysiology of delirium remains limited. We have previously modelled delirium in rodents by superimposing LPS-induced systemic inflammation in animals with pre-existing hippocampal and thalamic neurodegeneration in the ME7 prion disease model^11^, demonstrating that LPS impaired working memory in animals with prior degeneration, in a shallow water T-maze task, but did not do so in normal animals. Others demonstrated LPS impairments of working memory selectively in older mice^12,13^ and more recently we have shown that the vulnerability of aged mice to such deficits shows significant heterogeneity but is correlated with loss of integrity of hippocampal synapses and with white matter microgliosis and myelin disruption^14^.

Certain cytokines including IL-1β and IL-6 have mechanistic roles in driving sickness behaviour responses, but agents that trigger sickness behavior including LPS, IL-1β and TNFα all also reliably induce hypoglycemia^15,16^. Recent studies replicating LPS-induced hypoglycemia, demonstrated that the glycolysis inhibitor 2-deoxyglucose exacerbated the suppression of locomotor activity despite blocking IL-1 production and showed that exogenous glucose was sufficient to suppress the sickness behaviour response and mitigate the working memory impairments after LPS challenge^17^. In clinical FDG-PET studies it has been shown that delirium is associated with reduced glucose utilisation in several areas of the brain^18–20^ and CSF biomarker studies have indicated increased brain insulin resistance and elevated ketone bodies in hip-fracture patients with delirium^21^. Together these studies support a role for disrupted brain energy metabolism in delirium and, in particular, suggest that reduced or impaired glucose metabolism could disrupt circuit function that is vital to normal attention and cognition in delirium.

When blood glucose becomes scarce, the liver generates ketone bodies from fat, to provide an alternative energy source to sustain vital function and spare glucose utilisation in the heart, brain, kidneys and muscles^22^. β-hydroxybutyrate (BHB) is the most abundant, but acetoacetate and acetone also contribute as energy substrates. During stress responses in humans cortisol and catecholamines can promote the production of ketone bodies^23,24^ and experimental sepsis also induces ketogenesis in humans^23^ and in mice^25^. As well as its role as an alternative energy substrate, several signalling functions have been revealed for BHB: it has been shown to induce widespread transcriptional changes by inhibiting class I histone deacetylase, leading to reduced oxidative stress^26^ and also can inhibit the NLRP3 inflammasome^27^, an important mediator of LPS-induced inflammation. Interestingly, NLRP3 inhibitors^28^, HDAC inhibitors^29^ and ketogenic diets^30,31^ have all been shown to mitigate dementia– or age-related cognitive decline.

In the current study we hypothesized that LPS-induced systemic inflammation would incude widespread changes in energy metabolism. Among which we proposed that ketone bodies would be elevated by LPS and that oral ketone body treatment, with the ketone ester (KE) bis-hexanoyl (R)-1,3-butanediol (C6×2BD), would mitigate LPS-induced hypoactivity and acute cognitive dysfunction in aged mice. To that end we examined the effects of LPS (250µg/Kg) and C6X2BD (3g/kg) on blood energy metabolites, energy utilisation and on the production of pro-inflammatory cytokines. The real-time utilisation of different energy sources and expenditure of energy was quantified by the respiratory exchange ratio (RER) in indirect calorimetry (metabolic cage) experiments and, using radioisotope-labelled glucose and betahydroxybutyrate (βHB), we also monitored how LPS and ketone body administration affected the whole-body oxidation of blood glucose and ketones. Using a mass-spectrometry metabolomic approach we investigated the impact of these treatments on key energy metabolism pathways. Finally, we tested whether the oral administration of KE could alleviate LPS-induced delirium-like effects in a cognitive flexibility task. Our results show that oral KE mitigates brain energy disruption by LPS, reduces the sickness behaviour response and largely prevents delirium-like cognitive deficits in aged mice without altering the pro-inflammatory response.

## Materials and methods

### Animals

All animal experiments other than the metabolic measurements (metabolic cages and stable isotope labeling) were carried out at TCD in accordance with European Commission Directive 2010/63/EU and were performed following ethical approval by the TCD Animal Research Ethics Committee and licensing by the Health Products Regulatory Authority (HPRA). At TCD, C57BL/6J mice aged 20 to 25 months, equal numbers male and female, were obtained from the National Institute on Aging’s (NIA) Aged Rodent Colony, then housed at 21°C with a 12/12 h light/dark cycle (lights on 8 A.M. to 8 P.M.) with food and water available *ad libitum.* The metabolic measurement experiments were carried out at the Buck Institute, also using aged C57BL/6J mice obtained from the NIA Aged Rodent Colony. These experiments were conducted when the mice were 19 months old. Animals were kept in a specific pathogen-free barrier facility under a 12/12 h light/dark cycle (lights on 6 A.M. to 6 P.M.) with food and water available *ad libitum*. Mice were housed in groups of 4 or 5 until the day before the scheduled experiment, at which point they were single-housed with a hideout and nesting materials. These animals were maintained according to the National Institutes of Health guidelines, and all experimental procedures were approved by the Institutional Animal Care and Use Committee (IACUC) at the Buck Institute for Research on Aging.

### Treatments

Mice were treated intraperitoneally (i.p.) with bacterial endotoxin 250 µg/kg LPS (Salmonella Enterica, equine abortus, Sigma, L5886) or 10 ml/kg saline. Mice were dosed 1 hour later orally with either 3g/kg bis-hexanoyl (R)-1,3-butanediol (C6×2BD, **Fig. S1**), provided by Component Health) or 3ml/kg canola oil (Can) with flexible gavage needles (Instech). C6×2BD is a diester compound comprising two moieties of hexanoic acid ester-bound to a moiety of R-1,3-butanediol. When consumed, C6×2BD is rapidly hydrolyzed to its constituents in the gut^32^, which are all rapidly converted to ketone bodies in the liver. R-1,3– butanediol is converted directed to BHB by alcohol and aldehyde dehydrogenases, while hexanoic acid is a medium-chain fatty acid that is oxidized to generate acetyl-CoA and ketone bodies (BHB and acetoacetate) via the endogenous ketogenesis pathway regardless of nutritional state or insulin levels.

### Behavioural assays

#### Activity

To investigate spontaneous activity, distance travelled, rearing and grooming were observed in an open field. Briefly, mice were allowed to freely and individually explore an opaque open field arena (58 x 33 x 19 cm) for 5 minutes. Total distance travelled, time spent immobile and rearing were recorded: captured using a video camera and analysed using AnyMaze software (Version 4.99). Rearing is here defined as the mouse standing on its hind legs with its front paws not touching the floor, while its head is raised in an effort to survey its surroundings, displaying an exploratory behaviour rather than mere locomotor activity.

### Cognitive function: Learning, retention, flexibility

To investigate cognitive flexibility, a ‘paddling’ Y-maze visuospatial task was used, as previously described^33^. This task has been custom-designed to test cognitive function during acute sickness, is assessed on correct and incorrect responses only and is not confounded by reduced motivation or by decreased speed of movement. A clear perspex Y-maze consisting of three arms with dimensions 30 × 8 × 13 cm was mounted on a white plastic base. The distal end of each arm contained a hole, 2 cm above the floor and 4 cm in diameter, into which a black cylindrical escape tube or a blocked black cylinder could be inserted to allow, or to block, exit from that arm. On exiting through this escape tube mice willingly enter a black ‘burrowing’ tube in which they can escape the water and be returned to their home cage. At the outside of the maze each of the three plastic inserts were covered by burrowing tubes so that both from the outside and from the centre of the maze, all arms looked identical. As such, the maze is solved using extra-maze cues. The maze was filled with 2 cm of water at 20°C, sufficient to motivate mice to leave the maze by walking to an exit tube, from where they are returned to their home cage. Visual cues were placed at and around the end of each arm of the maze, one of them being the experimentor. Mice were placed in one of two possible start arms in a pseudorandomized sequence for 12 trials (on each training day) and the groups were counter-balanced with respect to the location of the exit and start arm. For any individual mouse the exit arm was fixed (to train and test reference memory). Mice were trained for 11 days in order to ensure that all mice could consistently find the exit at baseline. All mice were aged 22 ± 2 months. An arm entry was defined as entry of the whole body, excluding the tail, and a correct trial was defined as entry to the exit tube without entering other arms. After this training period mice were injected intraperitoneally with 250 µg/kg LPS and tested for retention of prior memory in this maze over 4 trials. KE or Canola oil were given by oral gavage at 1 hour post-LPS, as before. Each mouse started at 150 minutes post-LPS challenge. Then all mice had their exit moved to a different arm. The ability of treated mice to learn the new location of the exit was assessed over 16 trials starting from 3 hours after LPS injection. 12 further trials were performed the next day, starting at 24 hours after LPS. Number of arms entered, successful trials (mouse exiting the maze without entering any incorrect arms) and sequence of entries (arms 1, 2, 3) were recorded.

### Blood glucose and BHB measurements

For serial blood measurements, mice were placed in a plastic restraint tube, with the tail exposed. A tail vein was nicked with a 30-G needle facilitating the collection of a droplet of blood. Blood concentrations of glucose and BHB were assessed using point-of-care devices (Accu-Check® Aviva glucometer and Keto-Mojo GK+ reader). At 6 hours after LPS, blood glucose and BHB were measured from terminal blood collected immediately after the beginning of transcardial perfusion, following IP sodium pentobarbital injection.

### Plasma ELISA

Under terminal anaesthesia, blood was collected into heparinized tubes directly from the right atrium. Blood samples were then centrifuged at 1500 g for 15 minutes at 4°C. Plasma supernatants were then collected in a new tube and stored at –20°C until use. Plasma concentration of pro-inflammatory cytokines were measured using commercial kits, either R&D DuoSet (TNF-α: DY410; CCL2: DY479), Biolegend ELISA MAX (IL-6: 431301) or R&D Quantikine (IL-1β: MLB00C). Manufacturer’s instructions were followed and either duplicates or serial dilutions were made to increase accuracy. Plasma samples were used undiluted for IL-1β and TNF-α, while dilutions were used, ranging from 1/2 to 1/256 for CCL2, and from 1/2 to 1/128 for IL-6.

### RT-qPCR

6 hours after LPS challenge, mice were terminally anaesthetized with intraperitoneal sodium pentobarbital and transcardially perfused with heparinised saline. Brains were collected and hippocampi, cortices, and hypothalami were snap frozen in liquid nitrogen and stored at –80°C until used. Hippocampi halves were lysed in Trizol (Invitrogen) and RNA was extracted following the manufacturer’s instructions. An additional DNAse step was added to ensure elimination of DNA. The RNA yield and quality of each sample were quantified based on optical density using the “NanoDrop” ND-1000 UV–vis spectrophotometer (Thermo Fisher Scientific). cDNA synthesis was carried out using a High Capacity cDNA Reverse Transcriptase Kit (Applied Biosystems, Warrington, UK). Primers were either used with FAM/TAMRA probe with Faststart Universal Probe Master (Rox) (Roche) or without probe with Faststart Universal SYBR Green Master (Rox; Roche). Primers and probes were used at a final concentration of 200 nM in a reaction volume of 15 µL, and PCR were run in a Quantstudio 5 Real-Time PCR System, 384 well (Applied Biosystems). After 10 minutes at 95°C, cycling conditions were 95°C for 10 seconds, 60°C for 30 seconds, 50 cycles.

### Metabolic activity and tracer studies

Energy expenditure and the respiratory exchange ratio (RER) were measured using the Promethion Metabolic System (Sable, Nevada, US; https://www.sablesys.com/products/promethion-line/). Mice were first placed in individual housing and habituated overnight. On the following day, mice were transferred to single metabolic cages to acclimatize for 1-2 hours prior to LPS challenge to establish baseline measurements. Then, mice were treated intraperitoneally (i.p.) with bacterial endotoxin 250µg/kg LPS (Salmonella Enterica, equine abortus, Sigma, L5886) or 4ml/kg saline. Mice were dosed 1 hour later orally with either 3g/kg C6×2BD or 3ml/kg canola oil (Can) with flexible gavage needles and subsequently monitored in metabolic cages for 20 hours. The system continuously records the rate of carbon dioxide emission (vCO_2_) and rate of oxygen consumption (vO_2_) for each chamber. Mean RER, vCO_2_/vO_2_ reflects the percentage use of each substrate at the cellular level at any time: 0.7 for fats, 0.8 for proteins and 1.0 for carbohydrates. Mean energy expenditure (EE) in kcal/hr, calculated using the Weir equation^34^.

In additional experiments in the Promethion Metabolic system further mice were treated i.p. with 250µg/kg LPS or saline, dosed 1 hour later orally with either 3g/kg C6×2BD or 3ml/kg Canola with flexible gavage needles and, another hour later, were injected with either 12mg/kg of glucose or 8mg/kg of sodium D-3-hydroxybutyrate (BHB) tracer composed of heavy carbon (^13^C, Cambridge Isotope Laboratories, Inc., Tewksbury, MA, USA, Item No. CLM-1396-1 and CLM-3853 respectively). The tracer solution was prepared in Millipore water and administered at 8ml/kg. The dosages for both glucose and BHB tracers were previously tested to have minimal impact on the animals’ physiological blood glucose and BHB levels. For all treatment or tracer administrations, each mouse was briefly removed from the metabolic chamber as needed. For these analyses the chambers were connected to the Stable Isotope Gas Analyzer which allows the detection of tracer oxidation by the animal. Oxidation rates of the tracers were calculated by multiplying the metabolic rate (vCO2) to Δδ13C (raw tracer oxidation).

### Mass Spectrometry metabolomics

Primary metabolites including carbohydrates and sugar phosphates, amino acids, hydroxyl acids, free fatty acids, purines, pyrimidines, aromatics, and exposome-derived chemicals in snap-frozen hippocampi were analyzed by untargeted GC-TOF-MS (UC Davis West Coast Metabolomics Center, Davis, CA, USA). Metabolites were quantified by relative peak height but additional internal standards and calibration curves were utilized for the absolute concentrations of glucose, lactic acid, 3-hydroxybutyrate, citric acid, leucine, isoleucine, valine and tryptophan. Specifically, hippocampi samples extracted using 1mL of 3:3:2 ACN:IPA:H_2_O (v/v/v). Half of the sample was dried to completeness and then derivatized using 10 µL of 40 mg/mL of Methoxyamine in pyridine. Samples are shaken at 30°C for 1.5 hours. Then 91 uL of MSTFA + FAMEs to each sample and they are shaken at 37°C for 0.5 hours to finish derivatization. Samples are then vialed, capped, and injected onto the instrument. A 7890A GC coupled with a LECO TOF was used. 0.5 µL of derivatized sample is injected using a splitless method onto a RESTEK RTX-5SIL MS column with an Intergra-Guard at 275C with a helium flow of 1 mL/min. The GC oven is set to hold at 50°C for 1 min then ramp to 20°C/min to 330°C and then hold for 5 min. The transferline is set to 280°C while the EI ion source is set to 250°C. The Mass spectrometry parameters collect data from 85m/z to 500m/z at an acquisition rate of 17 spectra/sec.

Pooled QC samples and method blanks were injected every 10 samples alongside an external QC injection for instrument validation also injected every 10 samples. The untargeted portion was normalized by the average metabolite total ioncount (MTIC) of the samples and the relative peak height was used for statistical analyses. The targeted portion was normalized to the amount of sample on column using peak height. For the quantitative portion a 6-point curve is injected alongside the samples and used for concentration.

### Statistical analyses

Statistical analyses were performed in Graphpad Prism 10. Comparisons of pairs was performed using Mann-Whitney tests. Comparisons of multiple groups were performed using 2-way or 3-way (repeated measures) ANOVA or mixed-effects models where not all subjects had data at all time points. Where necessary, data were log-transformed to allow parametric statistics to be performed. Statistical analyses for specific experiments are detailed in the description of results or in the corresponding figure legend where appropriate. In metabolomics analysis, those metabolites for which absolute quantities were obtained were assessed for outliers using the ROUT test with Q set at 1%. Outliers were removed as follows: 1 outlier for both valine and leucine and 2 outliers for isoleucine and glucose. For those metabolites measured for relative abundance only, peak heights were first mean-centered, then divided by the standard deviation of each variable for scaling before any downstream analysis. Partial Least Squares Discriminant Analysis (PLS-DA) and volcano analysis of the metabolomic results were performed with all metabolites compared between LPS-treated and saline-treated animals, using the MetaboAnalyst web server (version 6.0, www.metaboanalyst.ca).

## Results

### LPS triggers sickness behaviour, hypoglycemia and ketosis in aged mice

We first assessed sickness behaviour, glucose and BHB following LPS challenge in C57BL/6J mice aged between 20 and 25 months. As expected, LPS decreased locomotor activity by more that 80% in aged mice (**Fig. 1A**, p<0.0001, Mann-Whitney U test). No significant differences between males and females were found in this experiment so data were pooled for analysis. Blood glucose was significantly and progressively decreased by LPS from baseline to 4, to 6 hours after injection compared to the saline group (**Fig. 1B**). Using mixed effects ANOVA, there was a main effect of time (F_1.4_, _43_=8.45, p=0.0023) and of treatment (F_1,31_=54.83, p<0.0001) on glucose levels as well as an interaction between time and treatment (F_2,60_=18.16, p<0.0001) with significant post-hoc differences at 4 and 6 hours (Bonferroni post-hoc tests, p<0.0001). For BHB, there was a main effect of time (F_1,31_=82.05, p<0.0001) and LPS had a bigger effect on BHB as time progressed (significant interaction: F_2,60_ 9.219, p=0.0003). In post-hoc comparisons, BHB was significantly higher in LPS than in saline at 6 hours (Bonferroni post-hoc test, p=0.0015).

**Figure 1:**
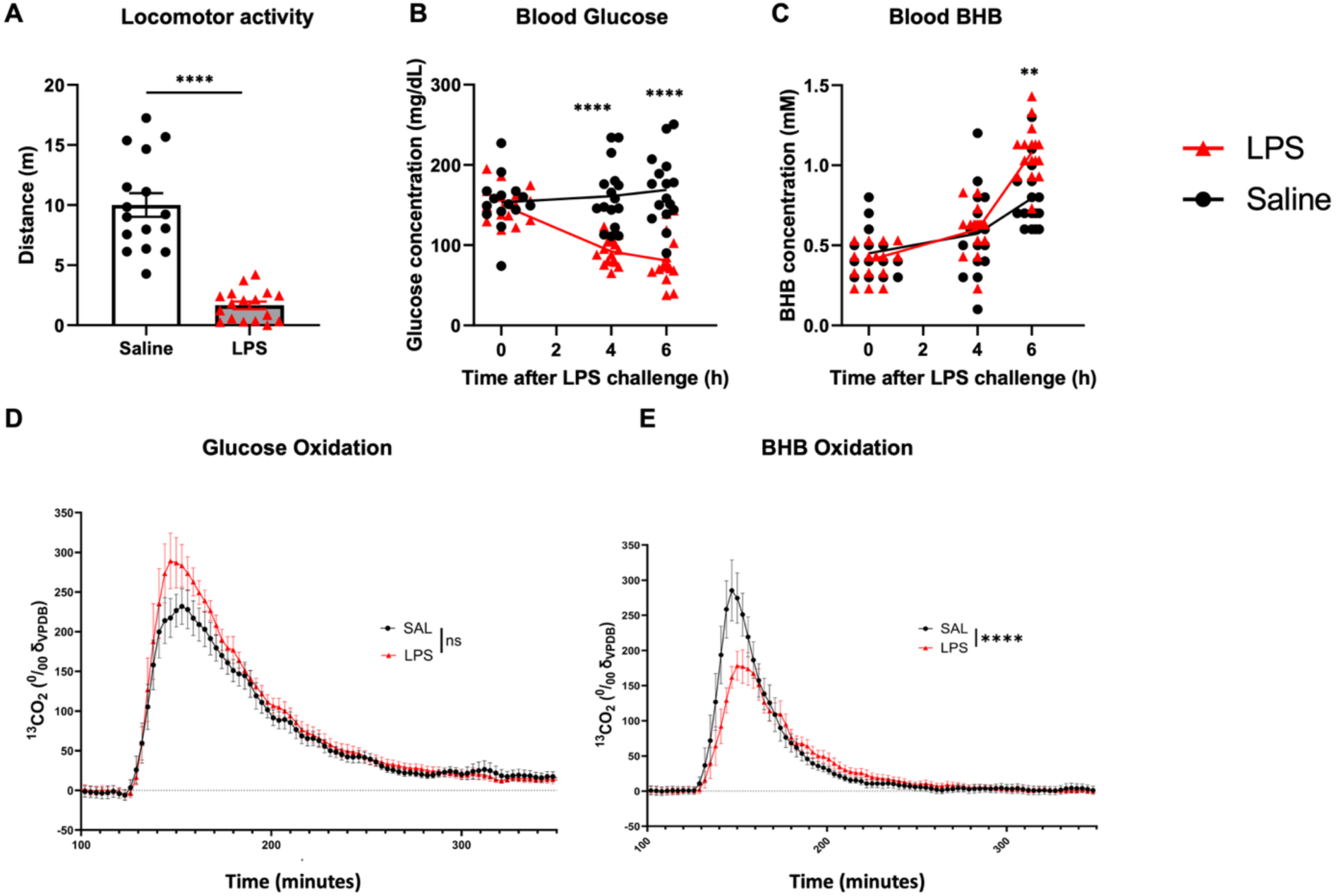
LPS induces locomotor suppression and altered glucose and ketone body metabolism. Aged mice (20-25 months) were injected with 250µg/kg LPS i.p. or 10mL/kg saline. **A)** Spontaneous activity was assessed 3 hours after LPS in an open field arena for 5 minutes. Data are shown as mean ± SEM, **** denotes p<0.0001 by Mann Whitney U test. Blood metabolites were assessed at baseline, 4 and 6 hours post-LPS using point-of-care devices to measure **B)** whole venous blood glucose (mg/dL) and **C)** whole venous blood BHB (mM). Blood metabolites data were assessed using mixed effects analysis, due to missing data on 1 time point, on 2 samples. Analysis showed significant decrease in glucose at 4 and 6 hours after LPS (fixed effect of LPS, p<0.0001, and significant post-hoc Bonferroni pair-wise comparison at both 4 and 6 hours, p<0.0001, denoted by ****). After a significant interaction of time and LPS, Blood BHB concentration was increased by 6 hours post-LPS by post-hoc Bonferroni comparison (p=0.0015, denoted by **). n=16 saline, n=17 LPS. **D)** Glucose catabolism rate measured as the derivative of the expired radiolabeled carbon dioxide following radiolabeled glucose injection 2 hours after LPS/saline treatment. **E)** BHB oxidation measured as the derivative of the expired radiolabelled carbon dioxide following radiolabelled BHB 2 hours after LPS/saline treatment. Experiment was balanced for sex; total n=7 saline and n=8 LPS.

Considering the evidence of hypoglycemia (Fig. 1B) and increased blood BHB (**Fig. 1C**) we interrogated whether LPS affects glucose and BHB oxidation. Two hours after i.p. injection with LPS mice were i.p. injected with either ^13^C glucose or^13^C BHB tracer and monitored in metabolic cages until 20 hours after LPS.^13^C exhaled in the form of CO_2_ can be detected almost immediately upon injection, indicative of tracer oxidation. We observed slightly more rapid glucose utilization in LPS-treated animals than in saline-treated animals, although this difference was not statistically significant (**Fig. 1D**): LPS x Time interaction in 2 way ANOVA: F_103,1339_=0.9180, p=0.7063 We also observed reduced oxidation of BHB during LPS-induced inflammation (**Fig. 1E**; LPS x Time interaction in 2 way ANOVA: F_86,1204_=3.019, p<0.0001) compared with that after saline treatment.

### Oral ketone ester alleviates LPS-induced hypoactivity

To assess whether elevated ketone bodies would mitigate the effects of LPS, mice were given 3g/kg KE or canola oil by oral gavage 1 hour post-LPS (250 µg/kg i.p.). LPS significantly reduced blood glucose (**Fig. 2B**) and KE treatment increased blood BHB (**Fig. 2C**). There were significant main effects of LPS (F_1,64_ =124.5, p<0.0001) and of time post-LPS (F_1.6,101_=36.19, p<0.0001) but KE did not have any significant effect on blood glucose with either LPS or saline. However there was a significant interaction between time and KE (F_2,126_ = 3.283, p=0.0407). Blood BHB levels were higher in KE treated mice by 4 hours, and LPS produced gradual but additive increases. Mixed-effects analysis revealed a main effect of KE (F_1,64_ =33.77, p<0.0001) and interactions between time and LPS (F_2,126_ =14.71, p<0.0001) and time and KE (F_2,126_ =11.47, p<0.0001). Post-hoc pairwise comparisons showed that LPS KE was significantly different to LPS CAN at 6h (p<0.004) and that Sal KE was already significantly different to Sal CAN by 4h (p=0.029).

**Figure 2:**
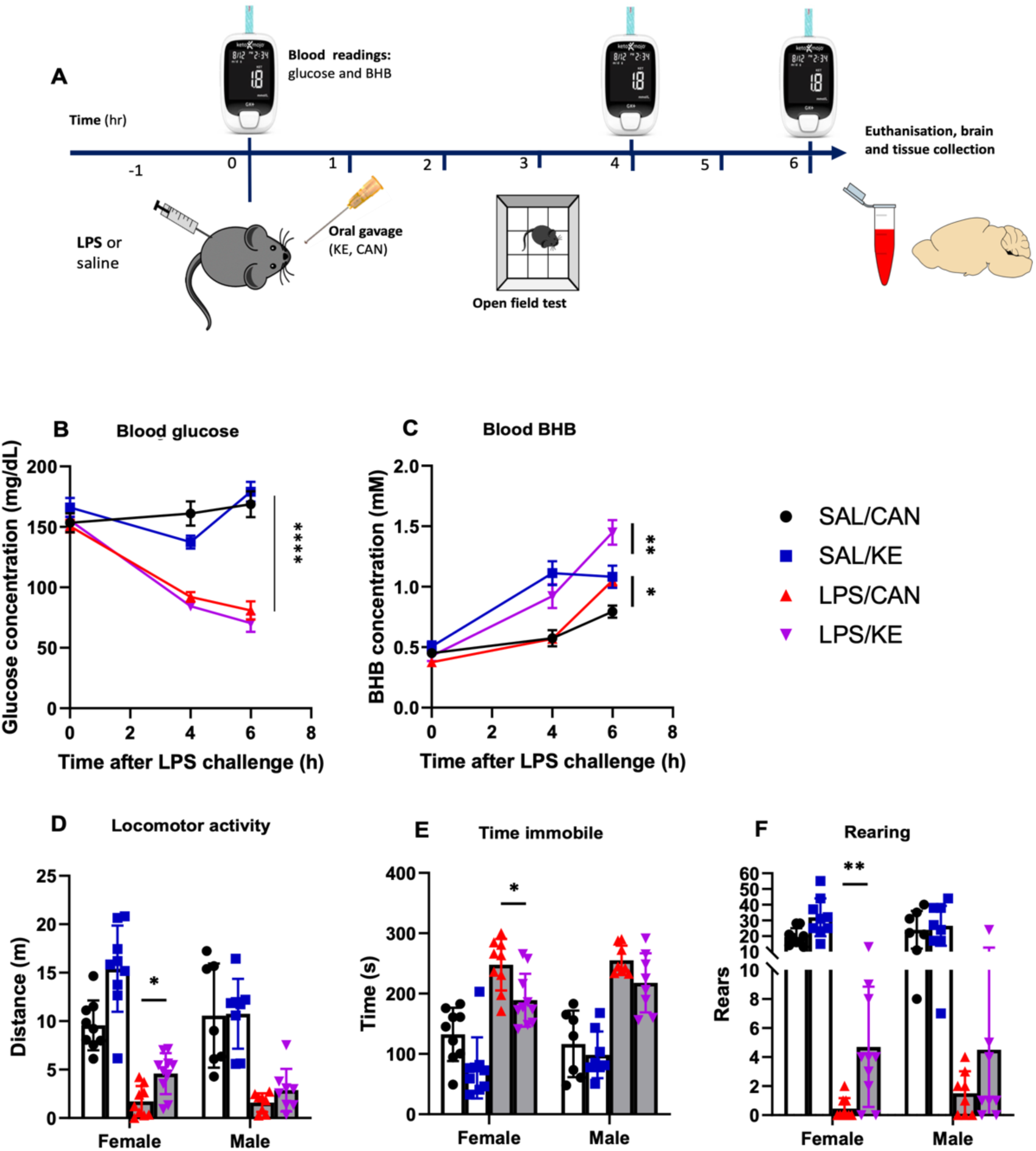
Oral ketone ester mitigates LPS-driven hypoactivity. (A) Experimental design. (B) Blood glucose (mg/dL) at baseline, 4 and 6 hours after i.p. LPS. Blood metabolites were analyzed using a mixed-effects model to account for a small number of missing values where blood collection was unsuccessful. **** denotes main effect of LPS; p<0.0001 as detailed in main text. C) Blood BHB concentration at baseline, 4 and 6 hours after i.p. LPS. * denotes post-hoc significant difference between Sal KE and Sal CAN at 4h, p=0.029, ** denotes post-hoc significant different between LPS KE and LPS CAN, p=0.004). (D) Locomotor activity, expressed as the total distance walked by mice in 5 minutes, (E) time spent immobile in 5 minutes in the open field arena and (F) number of rears in 5 minutes in the open field arena. Behaviour data were non-parametric and were log transformed in order to analyse using 3-way ANOVA with LPS, KE and sex as independent factors, followed by Bonferroni post-hoc comparisons. With the exception of baseline levels all Saline vs. LPS comparisons showed significant differences (p<0.0001) and are not denoted on graphs. Significant differences in Bonferroni post-hoc pairwise comparisons are denoted by * p< 0.05; ** p< 0.01. n = 9♀/7♂ SAL/CAN; 9♀/7♂ SAL/KE; 9♀/8♂ LPS/CAN; 10♀/8♂ LPS/KE. SAL: Saline; CAN: Canola oil; LPS: Lipopolysaccharide; KE: Ketone ester. Data are shown as mean±SEM.

In locomotor activity assessments (**Fig. 2D-F**) LPS administration significantly reduced distance travelled (F_1,60_ =61.38, p<0.0001), while KE increased it (F_1,60_ =13.5, p=0.0005). KE partially restored locomotor activity in females treated with LPS (p=0.0045), but not in males (interaction of KE with sex: F_1,60_ =4.024, p=0.049). Post-hoc pairwise comparisons showed that LPS KE was significantly different to LPS CAN (p=0.0045) while Sal KE was not significantly different to Sal CAN (p=0.378). LPS-induced hypoactivity was also reflected in the time spent immobile (Fig. 2E). LPS increased this (main effect of LPS F_1,60_ =120.3, p<0.0001) while KE decreased it (main effect of KE: F_1,60_ =15.38, p=0.002). Bonferroni post-hoc pairwise comparisons showed that KE significantly decreased the time LPS-injected female mice spent immobile (p=0.02) while not affecting this in males (p=0.768). The most striking effect of KE in LPS-treated animals was observed on rears (**Fig. 2F**). LPS strongly decreased the number of rears (F_1,60_ =163.8, p<0.0001) while KE increased rears. A main effect of KE was detected (F_1,60_ =9.898, p=0.0026) but the increased number of rears was significant in LPS-treated KE-treated females (Bonferroni post-hoc, p=0.008), but not in males (p=0.98).

### Peripheral and central pro-inflammatory cytokine production is not affected by oral KE

We proposed two non-exclusive theories to explain the impact of KE on LPS-induced hypoactivity: 1) based on reported inhibition of the NLRP3 inflammasome by ketones^27^, an early, acute treatment with KE may dampen the inflammatory response to i.p. LPS or 2) since hypoglycaemia contributes to hypoactivity in sickness behavior^17^, providing KE as an alternative energy source may alter energy metabolism to support spontaneous activity.

To test the first hypothesis, animals were euthanised at 6 hours post-challenge to evaluate blood and central cytokine production via ELISA and RT-qPCR respectively. Since the NLRP3 inflammasome is responsible for maturation and secretion of IL-1β we measured IL-1β release into plasma at 6 hours post-LPS. IL-1β was elevated by LPS (main effect of treatment: F_1,54_=29.25, P<0.0001) but there was no significant effect of the ketone treatment in either sex. Similarly, LPS significantly elevated levels of TNF-α (F_1,54_=21.65, p<0.0001), IL-6 (F_1,52_=11.89, p=0.0011) and CCL2 (F_1,52_=4.663, p<0.035) and these were also not affected by ketone ester treatment or by sex (**Fig. 3**). Hippocampal transcripts for *Il1b* (F_1,57_=25.45, P<0.0001) and *Tnfa* (F_1,57_=64.67, P<0.0001) were elevated in LPS-treated mice, while neither KE nor sex affected levels of these transcripts (Fig. 3B). In 3 way ANOVA analyses there were no other main effects or interactions. *Nlrp3* expression was increased in the hippocampus (F_1,57_=41.27, P<0.0001) and was higher in females (F_1,57_=47.64, P<0.0001) but this did not interact with LPS or KE. The expression of *Casp1* was not affected by LPS, ketone ester or sex. Thus, the ability of oral KE to mitigate the effect of LPS on activity is not the result of KE suppression of inflammatory cytokines.

**Figure 3:**
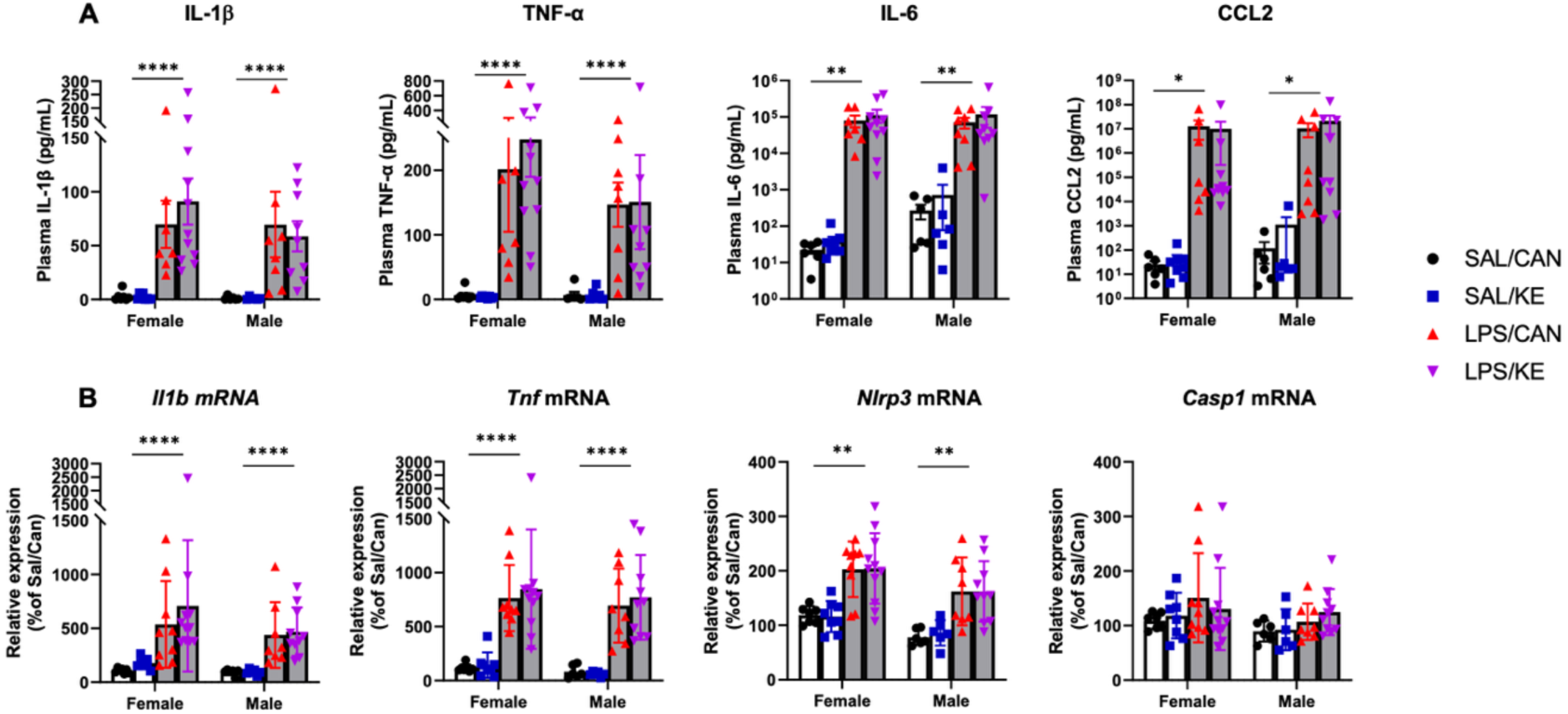
Central and peripheral pro-inflammatory cytokine production. Aged mice (20-25 months) were injected with 250µg/kg LPS i.p. or 10mL/kg saline, followed at 1 hour by KE or canola (3g/Kg by oral gavage. (A) Plasma cytokine concentrations were measured by ELISA. LPS significantly increased the plasma levels of IL-1β (F_1,54_=29.25, P<0.0001), TNF-α (F_1,54_=21.65, P<0.0001), IL-6 (F_1,54_=11.89, P=0.0011) and CCL2 (F_1,54_=4.663, P<0.0355). No main effect nor interaction were found with Sex nor KE. n= 6♀/6♂ SAL/CAN; 8♀/6♂ SAL/KE; 7♀/8♂ LPS/CAN; 10♀ /9♂ LPS/KE. (B) Hippocampal expression of transcripts for pro-inflammatory cytokines and inflammasome components were assessed by RT-qPCR. LPS significantly increased the expression of *Il1b* (F_1,57_=25.45, P<0.0001), *Tnfa* (F_1,57_=64.67, P<0.0001), *Nlrp3* (F_1,57_=47.64, P=0.0011) but not *Casp1*. Sex was also found to have a main effect on *Nlrp3* expression, were females had higher mRNA levels than males (F_1,57_=10.18, p=0.0023). No other main effect nor interaction was found. n= 7♀/6♂ SAL/CAN; 8♀/6♂ SAL/KE; 9♀/8♂ LPS/CAN; 11♀/10♂ LPS/KE. SAL: Saline; CAN: Canola oil; LPS: Lipopolysaccharide; KE: Ketone ester. Data shown as mean ±SEM. Significant main effects of LPS are denoted by *, ** and **** for p<0.05, 0.001 and 0.0001 respectively.

### Whole-body metabolism of LPS-treated animals switches to predominantly fatty acid oxidation

We pursued the second hypothesis, investigating whether whole-body energy metabolism and fuel type utilization are altered after LPS, and whether oral KE affects these changes. By measuring the respiratory exchange ratio (RER; the ratio of expired CO_2_ to consumed O_2_), the primary carbon source for energy metabolism can be estimated (fats vs. carbohydrates). Both male and female mice injected with saline showed a strong increase in RER during the dark period, indicating increased reliance on carbohydrate metabolism during the active phase (**Fig.4 A,B**). Conversely, animals injected with LPS see their RER fall to slightly above 0.7 (indicating increased fat oxidation) at 2 hours post-LPS and stabilize around that value for the remainder of the experiment. There was a main effect of LPS in females (2 way ANOVA: F_1,14_=23.24, p=0.0003) and in males (Mixed effects model due to some missing values: F_1,272_=63.49, p<0.0001). There was also an interaction of LPS and time for both females (2 way ANOVA: F_21,294_=12.46, p<0.0001) and males (Mixed effects model: F_21,,272_=23.32, p<0.0001). These data suggest that during systemic inflammation, fatty acids became the primary substrate for whole body oxidative metabolism.

**Figure 4.**
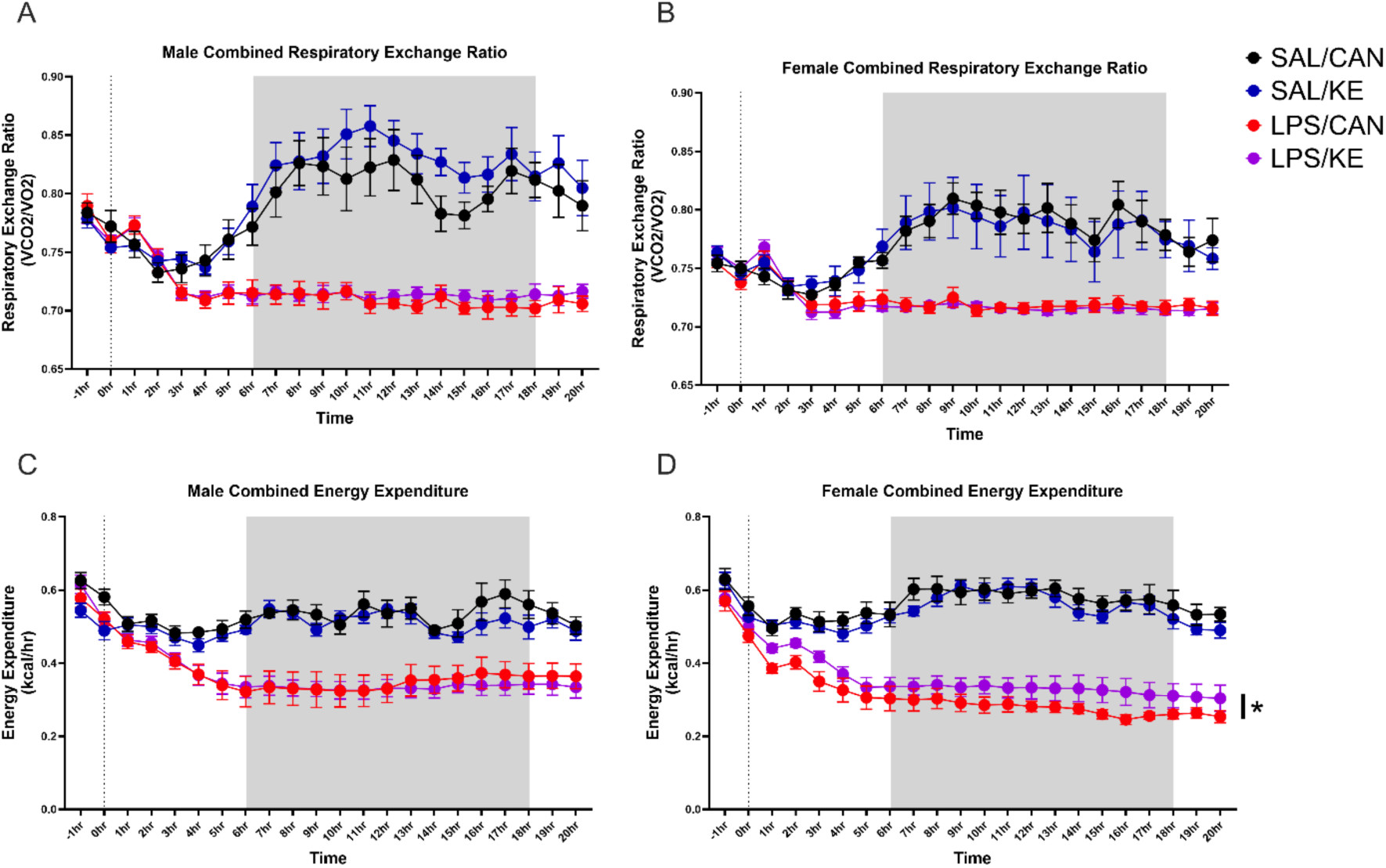
Animals were habituated in individual housing overnight and then in Promethion Metabolic cages for 1-2h, before injection with LPS (250µg/Kg), oral gavage with C6 ketone ester or canola oil 1 hour later and continued monitoring for a further 20 hours. Mean energy expenditure (kcal/hr) and the respiratory exchange ratio (RER) were calculated using measured expired CO_2_ and consumed O_2_. Main effects of LPS and of time are not denoted on the graphs. * denotes an interaction of LPS and C6 KE (F=4.76, p=0.047). n = 7 SAL/CAN, n = 8 each for SAL/KE, LPS/CAN, LPS/KE. This is applicable to both male and female experiments.

Mean energy expenditure (EE) in kcal/hr was also measured in the same apparatus (**Fig.4 C,D**), calculated using the Weir equation^34^. There was a very significant increase in activity starting just before the dark period and continuing throught that phase (main effect of time: F_21,294_=18.9, p<0.0001 in females and males: F_21,294_=20.01, p<0.0001). In animals treated with LPS this was very robustly suppressed in both males (main effect of LPS: F_1,272_=28.12, p<0.0001) and females (F_1,14_=112.3, p<0.0001). C6 KE very modestly increased energy expendure in female LPS-treated mice but not males (LPS x KE interaction in 2 way ANOVA: F_1,14_=4.76, p=0.047), indicating that modestly increased energy availability may be a mechanism by which KE affects behaviors in females.

### Impact of LPS and KE on brain energy metabolites

To assess brain energy substrate utilisation we performed mass spectrometry metabolomics on the hippocampi of those animals for which behavioural and inflammatory measures were presented above. Internal standards were analysed for absolute quantification for several pre-specified metabolites of importance in energy metabolism and insulin resistance (**Fig. 5a**). Despite robust peripheral hypoglycemia, LPS-treated mice did not show neuroglycopenia and brain and blood glucose were not significantly correlated (**fig. S2**). Treatment with KE produced a trend towards decreased glucose concentration. Conversely, treatment with KE resulted in elevated brain BHB in LPS-treated animals (post-hoc difference between LPS+KE and LPS+Can after significant interaction between LPS and KE in 2 way ANOVA: F_1,64_=4.2, p=0.043). Blood and brain BHB levels were correlated following LPS treatment (r = 0.51, p=0.002; **fig. S2**). The branched chain amino acids (BCAA), valine, leucine and isoleucine, markers of insulin resistance, were increased by LPS treatment (significant effect of KE treatment (F=5.56, 4.84, 5.47 respectively and p=0.02, 0.032, 0.23 respectively). There were no effects on the amino acid tryptophan. LPS significantly reduced levels of the glycolysis end-product lactate (main effect of LPS F_1,64_=16.7, p=0.0001) and significantly increased the citric acid cycle metabolite citrate (main effect of LPS F_1,64_=14.84, p=0.0003), but KE had no significant impact on either. Collectively these data indicate that the brain largely maintains levels of glucose, at least at 6 hours, despite systemic hypoglycemia, but lactate is nonetheless reduced. However all 3 BCAAs were increased by LPS, were strongly correlated with eachother (Pearson’s r ≥ 0.74), were moderately correlated with brain glucose concentration (**fig. 5b**; Pearson’s r ≥ 0.415) and the LPS-induced increases in all 3 were reversed by KE treatment. These data suggest brain insulin resistance and reduced brain glucose utilisation during endotoxemia and partial reversal of these by KE administration.

**Figure 5.**
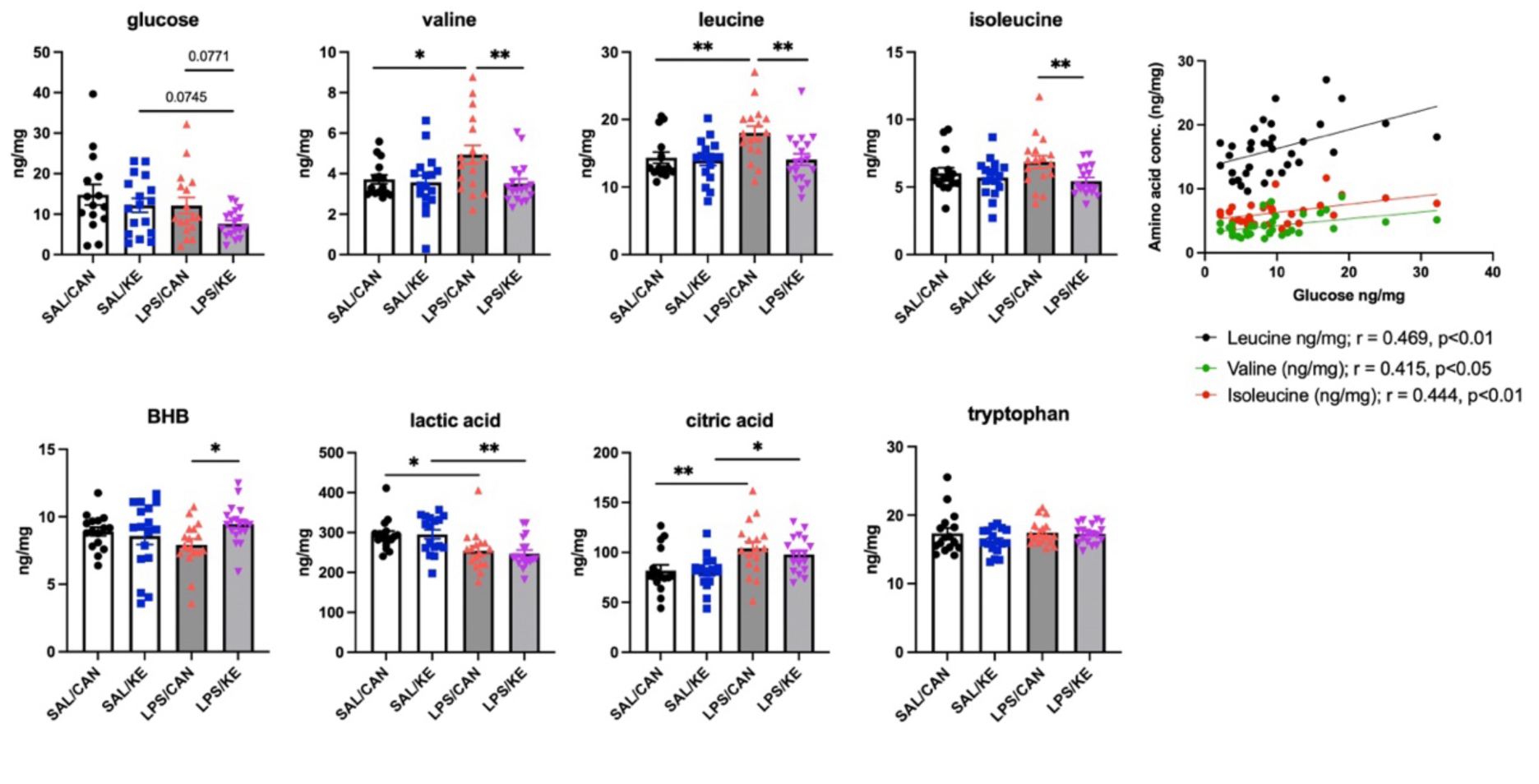
Prespecified metabolites analysed by Mass-Spectrometry. A limited panel was selected for absolute quantification based on the robust changes in peripheral glucose and ketone body metabolism following LPS. Aged mice (20-25 months) were injected with 250µg/kg LPS i.p. or 10mL/kg saline, followed at 1 hour by KE or canola (3g/Kg) by oral gavage and animals were terminally anaesthesised and perfused at 6h post-LPS. Samples arise from those animals assessed for sickness behaviour (in figure 2) and hippocampi were prepared for mass spectrometry, using internal standards to allow absolute quantification (a). *, ** denotes p<0.05 and 0.001 respectively using Fisher’s least significant difference tests for pre-specified comparisons after significant main effects or interactions using 2 way ANOVA. b) Analysis of correlation between glucose and each BCAA in all LPS-treated animals with calculated Pearson’s r and corresponding p value.

### Untargeted metabolomics to assess hippocampal metabolism during LPS-induced inflammation

Since treatment with KE altered brain glucose metabolism **(Fig. 5)** and increased spontaneous activity (**Fig. 2**), we proposed that untargeted mass spectometry metabolomics may reveal altered brain energy metabolism more broadly and in a manner that facilitates increased energy utilisation.

We first performed Partial Least Squares Discriminant analysis (PLS-DA) of all metabolites in LPS-treated versus saline-treated animals, revealing clear separation of these groups **(Fig. 6a).** Volcano plot of data from those animals showed that urea was the metabolite most significantly increased by LPS **(Fig. 6b,d)**. LPS can induce kidney injury and raised plasma urea and its elevation in the brain may reflect the convergence of LPS^35^ and age^36^ to produce increased blood brain barrier permeability^37^ to this molecule^38^. Glucose-6-phosphate (G6P), and the carbohydrates mannose and fructose, were the metabolites most substantially reduced by LPS **(Fig. 6 b,d),** while brain glucose itself is not reduced despite systemic hypoglycemia **(Fig. 5a, 6d)**. These carbohydrates were highly correlated with each other within individual animals (Spearman’s r = 0.68) and depletion of these carbohydrates accompanied sparing of hippocampal glucose **(Fig. 6d)**. Low mannose and fructose in particular were the glycolytic metabolites most strongly correlated with more severe sickness behaviour **(Fig. 6c)**, represented by suppression of distance moved and rearing behaviour and increased time immobile (Spearman r ≥ 0.47 for activity and r ≤ –0.45 for immobility). Those data are consistent with preservation of glucose levels and an LPS-induced switch to alternative fuels to glucose. BCAAs, particularly leucine, were also inversely correlated with activity measures (r=-0.48, –0.5 for distance and rears respectively). Consistent with entry of mannose and fructose into glycolysis, LPS also significantly increased transcription **(Fig. 6f)** of mRNA for enzymes accepting alternative substrates. These include HK2 (F_1,44_=29.99 p<0.001), capable of phosphorylation of mannose to M6P and aldolase B (F_1,44_=12.150 p<0.001), which converts fructose-1-phosphate to Dihydroxyacetone-phosphate (DHAP), allowing fructose to enter glycolysis more rapidly than glucose itself **(Fig. 6e)**. Further transcripts for metabolic enzymes and metabolite transporters are shown in **Fig. S3, S4**.

**Figure 6.**
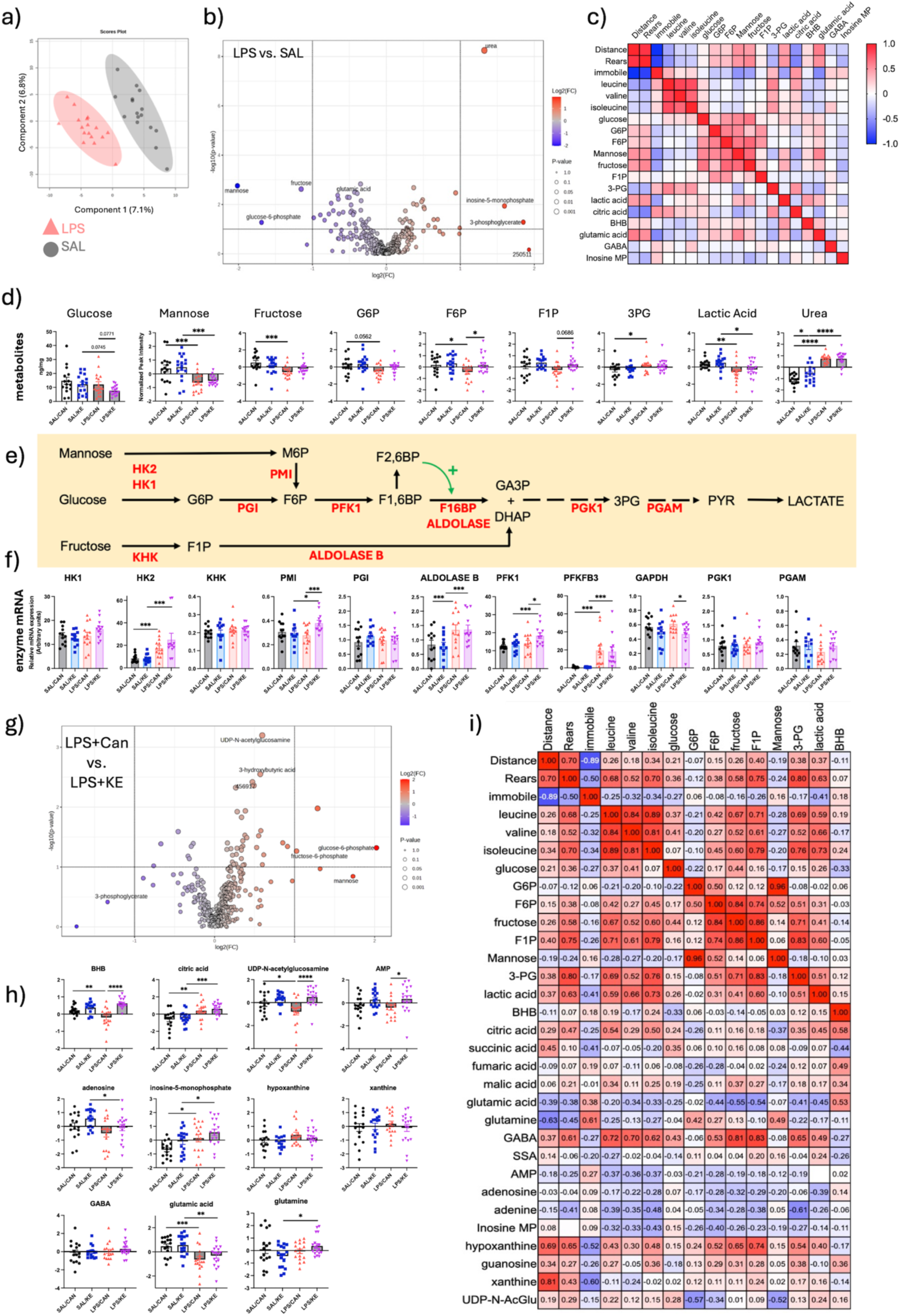
Untargeted metabolomic analysis of aged hippocampus after treatment with LPS ± KE. a) Principal component analysis of hippocampal metabolites 6h post-LPS (250 µg/Kg) versus saline-treated mice. b) Volcano plot of differentially detected metabolites in LPS versus saline-treated animals. c) Correlation matrix of sickness behaviour measures and key differentially detected and other glycolytic metabolites. d) Glucose concentration (absolute) and relative quantities of glycolytic metabolites in hippocampi from saline and LPS-treated mice, additionally treated with KE (3 g/Kg) or vehicle, calculated as normalised peak area. e) schematic of glycolysis showing metabolic intermediates and enzymes for both canonical and alternative carbohydrate substrates. f) Quantitative PCR for glycolytic enzymes in the opposite hippocampus from the same animals shown in panels 6a-d. g) Volcano plot of metabolites differentially detected at 6h in hippocampi from LPS-treated mice (250 µg/Kg) administered either KE (3g/Kg) or canola oil vehicle. h) Relative quantities of selected metabolites related to energy metabolism, energy status sensing and purine salvage in hippocampi from saline and LPS-treated mice, additionally treated with KE (3 g/Kg) or vehicle, calculated as normalised peak area. i) correlation matrix to examine the relationship between sickness behaviour measures and multiple metabolites in only LPS±KE animals. *, **, *** denote p<0.05, p<0.01 and p<0.001 respectively by Fisher’s least significant difference test (for normalised metabolites) and by Bonferroni post-hoc tests (for relative quantification of PCR), after significant effects in 2 way ANOVA analysis.

3-phosphoglycerate (3PG) was significantly increased in the hippocampus of LPS-treated mice **(Fig. 6b,d**; F_1,64_=6.38, p<0.05). *Pffi@3* was robustly induced by LPS (F_1,45_=98.434 p<0.001) and transcription of *Pffi1*, which is regulated by the PFKFB3 product F2,6BP, was also significantly induced by LPS (F_1,45_=6.585 p=0.014). However these were coupled with a significant decrease in lactate (main effect of LPS: F1,64=15, p<0.001) suggesting inefficient progress of glycolysis past the generation of 3PG, which would predict reduced hippocampal ATP generation. Although 3PG accumulation indicates that glycolysis progresses sufficiently to achieve the first ATP-generating reaction of glycolysis (PGK1->3PG) low lactate was one of the strongest predictors of inactivity in LPS-treated animals (r=0.47, 0.5 for distance and rears, **Fig.6c**). The strongest correlates of LPS-induced sickness were citric acid (negative correlation ≤0.52) and glutamic acid (positive correlation of ≤0.53). Thus, there are significant changes in glycolytic metabolites.

Several of the above changes trend towards a return to normal upon treatment with KE (**Fig.6g**). The volcano plot of LPS vs LPS+KE shows a reduction in abundance of 3PG that did not reach statistical significance. There were also significant increases in G6P and F6P. Mannose was also increased, though not quite statistically significant (**Fig. 6g**).

In 2 way ANOVA, with both LPS and KE as independent factors, F6P was significantly increased by the addition of KE (**Fig. 6d**) (p<0.05 by Fisher’s LSD). Transcription of phosphomannose isomerase (PMI), the enzyme that converts M6P to F6P for entry into glycolysis, though not increased by LPS, was significantly increased by KE treatment (interaction of treatment and intervention: F_1,45_=5.986 p=0.01). Transcription of *Pffi1* was also significantly increased by KE treatment compared to LPS+canola (p<0.05 post-hoc, after significant effect of LPS). Finally there was also significant decrease in transcription of GAPDH mRNA (p<0.05 Bonferroni post-hoc) although there was not a clear effect on 3PG levels.

Other metabolites relevant to brain energy status were altered by LPS and/or KE **(Fig. 6h)**. Importantly, BHB was depleted by LPS but robustly increased by KE treatment (main effect of KE, F1,58=23.88, p<0.0001 and significant post-hoc pairwise comparisons between LPS+Can and Saline: p<0.01; and between LPS+Can and LPS+KE: p<0.0001). Increased AMP typically indicates a decline in cellular ATP but LPS actually reduced levels of AMP possibly indicating energy sufficiency and consistent with that, citric acid, another indicator of energy sufficiency was robustly elevated by LPS **(Fig. 5a, 6h;** Main effect F_1,64_=24.43, P<0.0001). Conversely, UDP-N-acetylglucosamine, a positive indicator of glucose availability^39^, was significantly reduced by LPS and restored by KE (Main effect of KE, F_1,64_=19.02; p<0.0001). Inosine 5 monophosphate (IMP) was also increased by LPS treatment and further increased by KE (main effect of LPS: F_1,64_=11.49, p<0.01 and main effect of KE: F_1,64_=5.406, p<0.05). There were changes in several metabolites in the purine salvage pathway with similar increases observed after KE for AMP, adenosine and IMP and more variable levels of hypoxanthine and xanthine.

Finally, there were substantial changes in the glutamine/glutamate/GABA/succinate cycle. Glutamic acid was substantially decreased by LPS treatment (main effect of LPS, F_1,64_=21.25, p<0.0001) while glutamine was not significantly altered by LPS but was increased by KE treatment (interaction of LPS and KE, F_1,64_=4.27, p<0.05). GABA was not significantly altered by either treatment.

Considering the beneficial but variable effect of KE treatment on sickness behaviour **(Fig. 2**), we sought to assess which metabolites best correlate with behavioural improvements by examining correlations solely within the LPS+KE group. While concentrations of citrate, lactate and glutamate were strongly correlated with activity when analysing only LPS versus saline **(Fig. 6c)**, these metabolite alterations were not substantially reverted by KE **(Fig. 6d,h)** and correlations with activity in LPS+KE mice were weaker **(Fig. 6i)**. Nontheless, glutamate and, in particular, glutamine show significant negative correlation with activity (r=– 0.63 with distance) and GABA shows significant positive correlation (r = 0.61 for rears). GABA is negatively correlated with glutamate (r=-0.54) but is positively correlated with several of the glycolytic metabolites, which in turn are also positively correlated with activty in the open field (Lactate, F1P > 3PG > fructose).

Although AMP was not strongly correlated with activity there was a strong correlation between both xanthine and hypoxanthine and activity. Normalised peak areas were highly variable for these purines, across all experimental groups, and significant group differences were not evident, but their relative levels were the most strongly positively correlated with activity. KE-induced increase in AMP may indicate decreased metabolism of this purine or increased levels of ATP but how this contributes to hypoxanthine and xanthine levels and why these correlate with activity is not readily explained by the current data.

### Oral KE restores cognitive flexibility in LPS-treated aged C57BL/6J mice

Since brain energy metabolism changes with KE correlated with modest alleviation of sickness behaviors, we investigated whether KE would mitigate LPS-mediated deficits in cognition. We used a ‘paddling’ y-maze exit task^33^ to assess, in sequence, retention of consolidated reference memory and cognitive flexibility. We first trained animals to reliably find the exit without errors (11 days), using visuospatial cues (**Fig. 7a**). At 2.5 hours post-LPS or saline we tested retention and showed that animals reliably found the previously learnt location of the exit (**Fig. 7b**). Predictably, when the location of the exit arm was changed for each animal, the number of errors increased substantially in all animals. Thereafter the number of arms entered decreased over the first block of 4 trials in all groups, but LPS-treated animals began to make more errors than saline-treated or LPS+KE treated animals. Continued testing at 24 hours showed further improved performance in SAL+CAN and LPS+KE animals while LPS+CAN showed no improvement, remaining close to 50% correct (i.e. chance). Treatment statistically significantly affected cognitive flexibility (Mixed effects ANOVA, main effect of treatment, F_2,34_=6.514, P=0.004). Post-hoc analysis, using Bonferroni pairwise comparisons, showed that LPS+CAN mice performed significantly worse that LPS+KE on both blocks 6 and 7 (p=0.0035 and p=0.011 respectively) and worse than Sal+CAN on blocks 6 and 7 (p=0.049 and p=0.019 respectively). Thus LPS impairs the ability of aged mice to adopt a new strategy to exit the maze (cognitive flexibility) and treatment with KE, 1h post-LPS, protects against these deficits.

**Figure 7.**
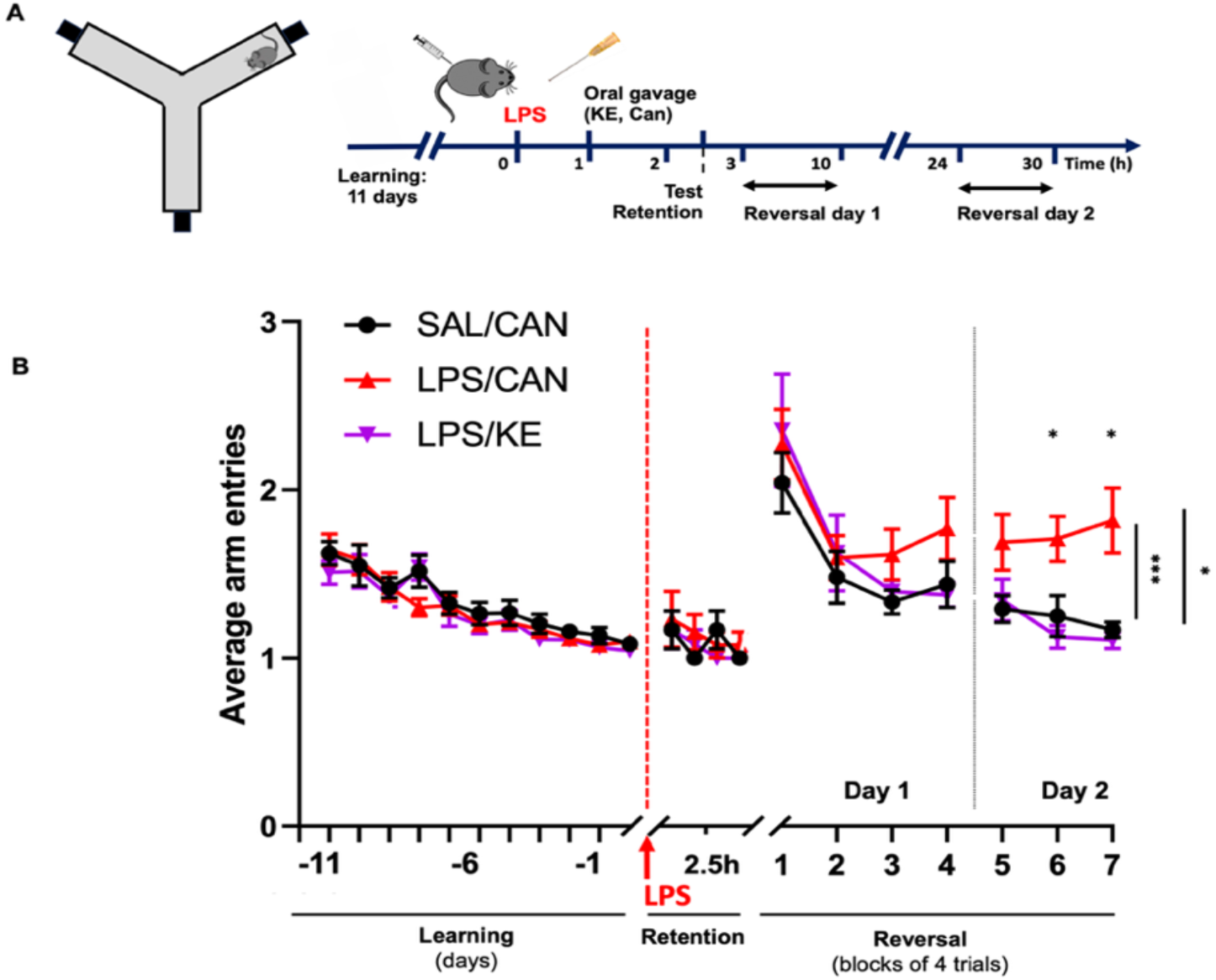
Oral KE prevents LPS-induced deficits in cognitive flexibility. (A) Experimental design. Animals were trained for 11 days to learn the location of the exit in this shallow wayer Y-maze. On day 12, mice were injected with LPS (250 µg/kg) or saline and given 3g/kg KE or canola oil 1h later. Retention of memory of each animal’s original exit was assessed through 4 trials at 2.5 hours after LPS, after which the exit was moved to a new arm and flexibility to locate the new exit was measured through 16 trials on day 1, and a further 12 trials day 2. (B) Errors were captured by measuring the number of arms entered before exiting the maze, where 1 arm entry is a successful trial. Treatment had a statistically significantly effect on cognitive flexibility (F_2,34_=6.514, P=0.004), and post-hoc comparisons found SAL/CAN and LPS/KE mice to perform better than LPS/CAN mice during the 6th block of 4 trials (p = 0.0489 and 0.0035, respectively) and on the 7th block (p = 0.0186 and 0.0112, respectively). SAL: Saline; CAN: Canola; KE: Ketone ester. n= 12 (6♀/6♂) for SAL/CAN; n=13 (7♀/6♂) for LPS/CAN, n=12 (6♀/6♂) for LPS/KE. Data are mean ±SEM.

## Discussion

In the current study we have demonstrated that LPS-induced systemic inflammation produces substantial alteration of whole-body energy metabolism with significant effects on brain energy metabolism. Within 4-6 hours of challenge with 250 µg/Kg LPS, we observed significant hypoglycemia as well as an increase in whole body FA utilisation, elevation of blood KBs and a slowing of peripheral KB oxidation. Despite this, treatment with exogenous ketone ester modestly protected against LPS suppression of spontaneous activity. Metabolomic analysis of the brain during LPS-induced hypoglycemia showed increased hippocampal concentrations of BCAAs, preserved brain glucose concentration and a prominent depletion of brain mannose and fructose. This shift away from glucose utilisation and towards insulin resistance was partially reversed by treatment with KE, suggesting that in the context of LPS-induced sickness KE may promote brain glucose utilisation and reduce the need for glycolytic alternatives. Consistent with this, KE treatment was sufficient to significantly protect against LPS-induced delirium-like deficits in cognitive flexibility. The data suggest that the significant reorganisation of energy metabolism required during acute peripheral inflammation may impose energetic constraints on the brain, which can compromise activity and cognition. Addressing the inflammation-induced metabolic reorganisation in the brain may be sufficient to prevent or reverse acute cognitive impairments occurring during systemic inflammation.

### Inflammatory versus metabolic effects of ketone bodies

Ketone bodies may exert both anti-inflammatory and metabolic effects in the brain. Although BHB can act as an anti-inflammatory molecule in inflammatory conditions via inhibition of the inflammasome^27^ we found no effect on circulating levels of IL-1β, TNFα or IL-6 in the blood nor brain transcription of *Il1b, Tnfa, Nlrp3* or *Casp1* following LPS. Previous studies report suppression of pro-inflammatory cytokines in LPS model of sepsis but doses of LPS were higher by a factor of 40 in that study and KE was administered more frequently^25^. Moreover, observed effects of KE on inflammation have been inconsistent in humans^40^. Here, at least under the conditions of moderate LPS-induced inflammation and during behaviourally relevant timescales, KE does not suppress measures of acute inflammation.

Thus we focussed on changes in energy metabolism. Initial findings with^13^C glucose and^13^C βHB showed that glucose tended to be more rapidly utilised during systemic inflammation and BHB, although also quickly utilised, was depleted more slowly in those with systemic inflammation.

Using indirect calorimetry to measure the RER showed that, at the whole body level, mice normally preferentially use carbohydrates but under systemic inflammation the RER falls to approximately 0.7, indicating increased FA oxidation, perhaps contributing to the endogenous generation of ketone bodies during LPS-induced inflammation **(fig. 1,2)**. Experimental endotoxemia in humans initially produces more rapid glucose utilisation, followed by insulin resistance and a slowing of glucose utilisation^41^. The importance of emergence of the ketogenic response is reflected by the observed necessity of reducing reliance on glucose utilisation for survival during sepsis^42^. In severe sepsis ketone bodies are generated from the oxidation of FAs and suppressing the ketogenic response via inhibition of PPARα activity exacerbates sepsis-induced lethality^42^. Thus, implementation of the ketogenic programme is adaptive during systemic inflammation and infection.

Although hypoglycemia, increased FA oxidation and ketogenesis all occured following LPS treatment in the current study, the spontaneous activity and energy consumption remained somewhat suppressed in all LPS-treated animals, even with KE treatment. This indicates that suppression of activity is not prevented simply by providing additional energy substrates. This is consistent with the hypothesis that behavioural suppression is regulated rather than reflecting an inability to source or expend energy. Nonetheless, treatment with exogenous ketone bodies did modestly lift the suppression of spontaneous activity in LPS-treated mice (fig. 2) and modestly increased energy expenditure in LPS-treated mice. Although this was only statistically significant in females, it was variable within both sexes and there were trends towards KE effects in males on all 3 measures of activity. Why effects might be greater in females is not clear. Differential impacts of KE on behaviour could relate to dimorphic behavioural responses to the inflammatory stimulus. LPS has been reported to have more robust effects on sickness behaviour in females than males^43^ although this was not apparent in LPS-treated males versus females in the current study and has also not been universally observed in the literature^44^. However, KE did modestly increase energy expenditure in indirect calorimetry experiments herein and might explain selective improvement in females.

### Altered brain energy metabolism during LPS-induced systemic inflammation

The very robust change observed in whole body metabolism during LPS-induced systemic inflammation indicates a shift away from peripheral glucose metabolism and mass spectrometry in hippocampal tissue indicated significantly altered brain glucose metabolism.

In an unbiased analysis of all metabolites covered in this metabolomic analysis, mannose, fructose and fructose 6 phosphate were among the most depleted metabolites in mice treated with LPS (**Fig. 6b**) and fructose-1-phosphate and glucose-6-phosphate were also somewhat depleted. All of these changes occur despite brain glucose concentration remaining stable during systemic hypoglycemia at 6 hours, when behavioural deficits are evident. Although the fold differences were relatively small, correlating spontaneous activity with metabolites relating to glycolysis, insulin resistance, citric acid cycle and excitation/inhibition balance, showed that reductions in mannose, G6P, fructose and F6P were correlated with spontaneous activity (**Fig. 6c**): lower levels of these carbohydrates predict lower activity. Conversely, glucose showed a very weak correlation with activity while decreased lactate, decreased glutamic acid and increased citric acid also showed strong correlations with activity when LPS was compared with saline.

The observed changes in these metabolites, at a single timepoint, cannot inform on glycolytic flux, but further support is offered by changes in expression of key enzymes (**Fig. 6f**). There was clear upregulation of HK2, which can also accept mannose as substrate and of aldolase B, which is the enzyme that converts F1P to DHAP and facilitates entry into glycolysis. While mannose can enter the cell via glucose transporters, it frequently accumulates as M6P and may inhibit glycolysis, as has been shown in multiple cancers^45^. However the increased expression of PMI, which was induced by KE here, can convert it to F6P and thus facilitate its contribution to the maintenance of glycolysis^46^.

Mannose has been shown to be used in glycolysis and in volunteer studies it has been shown to be the most successful substrate at compensating for glucose and preserving normal brain function in experimental hypoglycemia^47^. Uptake across the blood brain barrier, transport into neurons via GLUT transporters, phosphorylation by hexokinase and conversion to F6P by PMI have all been demonstrated^48–50^. PMI is also active in astrocytes^51^. Fructose has also been shown to be used by the brain^52^, feeding into glycolysis, via F1P and DHAP. Therefore this also potentially provides a short-term source of carbons for glycolysis. However F1P does join glycolysis below the key rate limiting step, phosphofructokinase 1, so it may operate unregulated and could lead to depletion of ATP and increased levels of uric acid^53^.

Collectively these findings may suggest that, with peripheral pressure on glucose concentration, the brain begins to deplete mannose and even fructose, presumably through use in this alternative glycolytic pathway. There is an earlier literature to support such a strategy^47^. In doing so, glucose levels in the brain may be preserved. We demonsrate that LPS increases brain concentrations of branched chain amino acids, valine, leucine and isoleucine, well established markers of insulin resistance^54^, that glucose levels are maintained and that KE treatment reverses the BCAA changes and produces a proportionate depletion of glucose. These data suggest development of brain insulin resistance that is reversible by ketone bodies.

In addition, LPS robustly increased transcripts for PFKFB3, which is known to promote activity of PFK1 and to increase glycolytic flux^55^. This might be beneficial in numerous cell types, including activated macrophages or microglia, or in particular metabolic scenarios where glycolysis may otherwise become suppressed. PFKFB3 activation can potentiate insulin-induced GLUT4 translocation to the plasma membrane and improve glucose uptake^56^ and we also observed *Glut4* expression to be increased by LPS (**Fig. S3**). However excessive activation of *Pffi@3* in neurons can be detrimental to neuronal integrity and cognition^57^. The latter findings are reported in the context of glucose sufficiency, however during hypoglycemia, upregulation of *Pffi@3* may constitute a compensatory mechanism. In this context it is also important to be clear that neither the metabolomic nor transcriptional analyses here are cell type-specific, so these changes may be occurring in any of a number of cell types and this requires further investigation. Transcriptional expression of glucose and ketone body transporters indicates LPS-induced increases in monocarboxylate transporters (MCT) 1 and 4 as well as glucose transporter 4 (**Fig. S4**). MCTs 1 and 4 are thought to be predominantly astrocyte-expressed (Pierre & Pellerin, 2005) while GLUT4 is predominantly neuronal and insulin-dependent.

Perhaps consistent with the utilisation of alternative carbohydrate sources to maintain glycolysis during acute inflammation is the observation that 3PG was increased in the hippocampus by LPS, ensuring that at least some hippocampal ATP is generated by glycolysis during systemic hypoglycemia. It is difficult to interpret whether this implies an abundance of 3PG during LPS-induced sickness because it cannot be used or whether this is a regulated preservation of 3PG levels. However it is striking that citric acid is also increased by LPS treatment and remains increased after KE treatment, suggesting that the citric acid cycle remains replete with either pyruvate or BHB, arguing for a regulation of, rather than a failure of, energy metabolism. Levels of both 3PG and citrate remain correlated with spontaneous activity in LPS+KE mice.

### Cellular sensors of nutrient and energy status

Given the systemic hypoglycemia, simultaneous with normal brain glucose and the subsequent provision of additional fuel in the form of KE, questions about the status of molecules that signal energy status arise. UDP-N-AcetylGlucosamine (UDP-GlcNAc) is produced from G6P via the hexosamine biosynthesis pathway and is an important precursor for glycosylation of proteins crucial for adaptation of neurons to increased ATP demand^58^. High levels of UDP-GlcNAc indicate abundant G6P, therefore nutrient sufficiency. The low levels observed following LPS may indicate hippocampal detection of some limitation in nutrient supply but the observed clear increase after KE treatment would appear to indicate nutrient sufficiency in that scenario (**Fig. 6h**), consistent with a role of BHB as an available brain fuel in this experimental model.

AMP levels typically rise when energy balance is low, activating AMP-activated kinase (AMPK) which can alter both brain and whole body metabolism to promote fatty acid oxidation and maintain cellular energy levels^59,60^. Previous studies with i.p. LPS reported decreased AMPK activity in both the hippocampus^61^ and in the hypothalamus^62^. The latter triggers suppression of glucogenesis in the liver, contributing to LPS-induced hypoglyemia. In the current study, hippocampal AMP levels were not increased by LPS, arguing against reduced brain ATP levels during LPS-induced illness. However AMP levels were increased by KE treatment. This may indicate that BHB facilitates increased usage of ATP. Consistent with that, there is tentative evidence of activated purine salvage pathways with increased inosine monophosphate and positive correlations between xanthine, hypoxanthine and spontaneous activation. However these are speculative explanations and further studies are required to substantiate these possibilities.

### Ketone bodies, brain energy metabolism and functional deficits

Although ketone body treatment does increase brain BHB concentration (**Figs. 5, 6h**), the BHB level itself does not correlate with activity. Thus, one hypothesis arising from the current study is that sharply declining levels of circulating blood glucose drives the implementation of strategies for the brain to mobilise, and use, alternative fuels. Providing alternative fuels such as KBs may lessen the pressure on the brain to implement the mannose/fructose strategy and thus tends towards normalising their levels but KBs themselves may be turned over rather quickly and therefore may not be a prominent correlate of activity even if important in facilitating it. Although depletion of alternative substrates may have contributed to the maintenance of brain glucose levels, the increase in BHB may have facilitated use of glucose and, in the early time points examined here, facilitated spontaneous activity and cognitive flexibility.

Although our blood analysis showed an elevation of blood BHB following KE gavage, our results do not capture the peak in blood BHB, which is reported to occur at about 30 minutes after gavage before returning to normal over several hours^63^. However, the BHB elevation is observable at 4 hours post-gavage and then continues to climb in those animals treated with LPS, perhaps facilitating additive effects of induced and administered ketone bodies. An improvement in cognitive flexibility was already apparent at 6 hours post-LPS but also continues on the following day. This raises the possibility that the benefit does not arise only from the consumption of ketone bodies as an energy source, but perhaps additional effects of the KE on energy metabolism more broadly. The most obvious explanation based on the data in the current study is the reversal of the LPS-induced increase in BCAAs, an accepted proxy for insulin resistance^64^. Reduced brain glucose utilisation has been reported in a number of studies related to sepsis^65,66^ and whether this is related to insulin resistance is not known.

It is of interest that KE treatment only modestly increased spontaneous activity and that BHB was more slowly oxidised in LPS-treated animals, consistent with the idea that efficiency of use of ketone bodies is subject to interindividual variability. In the current study the protective effect was more obvious in females than males and in metabolic cage experiments females also showed slightly more energy utilisation during ketone ester treatment than did LPS animals treated with canola control. Here it is important to consider that activity in both the metabolic cage and open field experiments is spontaneous and is not rewarded. By contrast, when tested in the shallow water Y-maze, in which there is a significant benefit to expending energy to escape this aversive environment, mice from both sexes benefited from the KE treatment. Although speculative, this may illustrate the cost:benefit nature of energy utilisation during acute illness. In the absence of a strong motivation to move and expend energy, sick animals expend minimal energy, but when there is a benefit to continued movement, animals can and do expend energy to reach a place of greater safety. This is consistent with the reorganisation of priorities concept of sickness behaviour^1,67,68^ and indeed provides indirect evidence that energy budget is a strong determinant of behavioural activity under circumstances of acute illness. The hypothalamus is highly activated by inflammation and, despite relatively widespread hypometabolism in LPS/sepsis studies^65,66^, glucose uptake was increased in the hypothalamus^42^. In fact the ventromedial hypothalamus contains glucose sensing neurons that are specifically activated by low glucose concentration, with roles in inducing counter-regulatory mechanisms in glucose metabolism^69^. Therefore the hypoglycemic state present in the current study is likely detected by these glucose sensing regions within hours of LPS challenge and their role in implementation of altered behaviour, and the impact of KE treatment in modulating that both require investigation.

### Ketone bodies and delirium

Ketone bodies are well described as an alternative energy source for brain energy metabolism and have been shown to beneficial in cognitive aging^30^ and in neurodegenerative diseases^70–72^. There is a growing interest in the idea that impaired glucose metabolism underpins cognitive dysfunction in dementias and other neurodegenerative diseases^73,74^ and there are both human and animal model data to support the idea that acutely interrupted glucose metabolism may be a key event in delirium: Insufficient glucose supply, triggered by LPS or by insulin, was causative of working memory dysfunction in an animal model of delirium superimposed on dementia^17^ and FDG-PET studies showed that reduced glucose uptake was associated with delirium in mixed medical patients^75^. Further to this, CSF studies in hip fracture patients have shown elevated BCAA levels consistent with insulin resistance and a robust increase of both BHB and acetoacetate in those with delirium^21^. Although those data come with the important caveat that those levels are measured at a single timepoint, usually within the first 24 hours post-fracture and pre-surgery, their elevation leads to 3 opposing hypotheses: 1) Elevated ketone body levels may be detrimental to cognitive function in those patients, 2) their levels may be elevated because of acute inflammation and those patients with the highest levels may be those most perturbed by the acute inflammatory insult or 3) their levels may be appropriately mobilised by the liver/blood but the brain may be impaired in its utilisation of the available BHB and AcAc. It is counterintuitive that a stronger supply of a usable energy substrate would be detrimental to maintaining normal brain function. Our current data indicate that treatment with ketone ester during acute systemic inflammation protects against deficits in cognitive flexibility.

While the current data likely refute the first hypothesis, they do not provide direct evidence to distinguish the second and third hypothesis. That is to say, it is not clear why patients with delirium have higher ketone body levels. An analogy might be drawn to a different disease state, heart failure, where elevated blood levels of ketone bodies were initially identified as a poor prognostic biomarker in patients while subsequent mechanistic experiments in rodents and humans determined that the use of ketone bodies by the failing heart was compensatory and beneficial to function^76^. The current study shows that systemic inflammation triggered by LPS is sufficient to drive elevation of BHB in the blood, so it is reasonable to suggest that patients with systemic inflammation do increase their KB synthesis. Although there is evidence from FDG-PET studies that patients with AD can still use KBs^77^, reduced ability to use ketones has been reported in aged mice^78^ and here, using^13^C labelled BHB, we show that animals treated with LPS are less efficient in their oxidation of BHB than are saline-treated animals. Even though glucose is limiting and ketone bodies are elevated, these aged animals, at least at the whole-body level, are not very efficient in using ketone bodies. Thus while elevated KE in human CSF might therefore be a marker of higher susceptibility to post-operative delirium^21^, our results in this reductionist model system, show that supplying additional ketone bodies is protective against inflammation-induced acute cognitive dysfunction. These relationships clearly require further study.

### Conclusion/Outlook

The current metabolic data broaden the knowledge base on the relationship between altered brain energy metabolism during LPS-induced systemic inflammation and it’s impact on behaviour and cognition. The brain’s use of alternative carbohydrates during inflammation-induced brain metabolism changes and the acute and reversible brain insulin resistance indicate significant brain adaptations to limiting peripheral glucose supply. The impact of KE on BCAAs in the current study suggest that ketone bodies may be useful in circumventing the impact of insulin resistance on normal brain function. While acknowledging the gap between the current animal model and clinical delirium, our data suggest that it could be of interest to test the ability of ketone bodies to mitigate delirium in patients with inflammation-induced delirium. Both PET imaging and fluid biomarker studies suggest changes in insulin resistance and glucose utilisation in patients with delirium and recent clinical trials offer tentative support for the idea of treatment with intra-nasal insulin^20^. The possible role of acutely induced insulin resistance in regulating behaviour and energy expenditure merits further investigation.

## Author contributions

PLH – performed experiments, analysed data, co-wrote manuscript. Designated first-named author on the basis of leading early contribution in both the findings and direction of the study.

MKKC – performed experiments, analysed data, co-wrote manuscript

JCM – performed experiments, analysed data

PD – performed experiments

HD – performed experiments

JCN – obtained funding, designed and analysed experiments and co-wrote the manuscript

CC – designed experiments, analysed data and co-wrote the paper

## Funding

This work was funded under a National Institutes of Health R01 (AG068025) to JCN and additionally by an IRC Laureate Award to CC (IRCLA/2023/1966).

## Conflict of Interest

JCN is co-founder, holds equity and is inventor on patents licensed to BHB Therapeutics Ltd., Selah Therapeutics Ltd. and BOPZ involving ketone esters. BHB Therapeutics provided the ketone ester used in these experiments but had no role in the design, execution or analysis of the experiments nor in the writing of the manuscript.

## Notes

### Summary of Updates

This is a minor revision to add citation of a number of articles on blood brain barrier permeability on page 15

## References

1. Dantzer, R. Cytokine, Sickness Behavior, and Depression. Immunol Allergy Clin North Am 29, 247– 264 (2009).

2. Draper, A. et al. Effort but not Reward Sensitivity is Altered by Acute Sickness Induced by Experimental Endotoxemia in Humans. Neuropsychopharmacology 43, 1107–1118 (2018).

3. Cunningham, C., Campion, S., Teeling, J., Felton, L. & Perry, V. H. The sickness behaviour and CNS inflammatory mediator profile induced by systemic challenge of mice with synthetic double-stranded RNA (poly I:C). Brain Behav Immun 21, 490–502 (2007).

4. Bluthé, R. M. et al. Role of interleukin-1beta and tumour necrosis factor-alpha in lipopolysaccharide-induced sickness behaviour: a study with interleukin-1 type I receptor-deficient mice. Eur J Neurosci 12, 4447–4456 (2000).

5. Palin, K. et al. Tumor necrosis factor-α-induced sickness behavior is impaired by central administration of an inhibitor of c-jun N-terminal kinase. Psychopharmacology 197, 629–635 (2008).

6. Godbout, J. P. et al. Exaggerated neuroinflammation and sickness behavior in aged mice following activation of the peripheral innate immune system. FASEB J 19, 1329–1331 (2005).

7. Elie, M., Cole, M. G., Primeau, F. J. & Bellavance, F. Delirium Risk Factors in Elderly Hospitalized Patients. J Gen Intern Med 13, 204–212 (1998).

8. Wilson, J. E. et al. Delirium. Nat Rev Dis Primers 6, 1–26 (2020).

9. Inouye, S. K., Westendorp, R. G. J. & Saczynski, J. S. Delirium in elderly people. Lancet 383, 911–922 (2014).

10. Davis, D. H. J. et al. Worsening cognitive impairment and neurodegenerative pathology progressively increase risk for delirium. Am J Geriatr Psychiatry 23, 403–415 (2015).

11. Murray, C. et al. Systemic inflammation induces acute working memory deficits in the primed brain: relevance for delirium. Neurobiol Aging 33, 603–616.e3 (2012).

12. Chen, J. et al. Neuroinflammation and disruption in working memory in aged mice after acute stimulation of the peripheral innate immune system. *Brain*, Behavior, and Immunity 22, 301–311 (2008).

13. Cunningham, C. & MacLullich, A. M. At the extreme end of the psychoneuroimmunological spectrum: Delirium as a maladaptive sickness behaviour response. Brain Behav Immun 28, 1–13 (2013).

14. Healy, D. et al. Susceptibility to acute cognitive dysfunction in aged mice is underpinned by reduced white matter integrity and microgliosis. Commun Biol 7, 1–15 (2024).

15. Rey, A. del et al. IL-1 resets glucose homeostasis at central levels. Proceedings of the National Academy of Sciences of the United States of America 103, 16039 (2006).

16. Evans, D. A., Jacobs, D. O. & Wilmore, D. W. Tumor necrosis factor enhances glucose uptake by peripheral tissues. *American Journal of Physiology-Regulatory*, Integrative and Comparative Physiology 257, R1182–R1189 (1989).

17. Kealy, J. et al. Acute Inflammation Alters Brain Energy Metabolism in Mice and Humans: Role in Suppressed Spontaneous Activity, Impaired Cognition, and Delirium. J Neurosci 40, 5681–5696 (2020).

18. Haggstrom, L. R., Nelson, J. A., Wegner, E. A. & Caplan, G. A. 2-18F-fluoro-2-deoxyglucose positron emission tomography in delirium. J Cereb Blood Flow Metab 37, 3556–3567 (2017).

19. Croteau, E. et al. A cross-sectional comparison of brain glucose and ketone metabolism in cognitively healthy older adults, mild cognitive impairment and early Alzheimer’s disease. Experimental Gerontology 107, 18–26 (2018).

20. Nitchingham, A. et al. Regional cerebral hypometabolism on 18F-FDG PET/CT scan in delirium is independent of acute illness and dementia. Alzheimer’s & Dementia 19, 97–106 (2023).

21. Titlestad, I. et al. Impaired glucose utilization in the brain of patients with delirium following hip fracture. Brain 147, 215–223 (2024).

22. Robinson, A. M. & Williamson, D. H. Physiological roles of ketone bodies as substrates and signals in mammalian tissues. Physiological Reviews 60, 143–187 (1980).

23. Vakharia, K. & Hinson, J. P. Lipopolysaccharide Directly Stimulates Cortisol Secretion by Human Adrenal Cells by a Cyclooxygenase-Dependent Mechanism. Endocrinology 146, 1398–1402 (2005).

24. Bahnsen, M. et al. Mechanisms of catecholamine effects on ketogenesis. American Journal of Physiology-Endocrinology and Metabolism 247, E173–E180 (1984).

25. Soni, S. et al. Exogenous ketone ester administration attenuates systemic inflammation and reduces organ damage in a lipopolysaccharide model of sepsis. Biochimica et Biophysica Acta (BBA) – Molecular Basis of Disease 1868, 166507 (2022).

26. Shimazu, T. et al. Suppression of oxidative stress by β-hydroxybutyrate, an endogenous histone deacetylase inhibitor. Science 339, 211–214 (2013).

27. Youm, Y.-H. et al. The ketone metabolite β-hydroxybutyrate blocks NLRP3 inflammasome-mediated inflammatory disease. Nat Med 21, 263–269 (2015).

28. Heneka, M. T. et al. NLRP3 is activated in Alzheimeŕs disease and contributes to pathology in APP/PS1 mice. Nature 493, 674–678 (2013).

29. Penney, J. & Tsai, L.-H. Histone deacetylases in memory and cognition. Science Signaling 7, re12– re12 (2014).

30. Newman, J. C. et al. Ketogenic Diet Reduces Midlife Mortality and Improves Memory in Aging Mice. Cell Metab 26, 547–557.e8 (2017).

31. Zhou, Z., Kim, K., Ramsey, J. J. & Rutkowsky, J. M. Ketogenic diets initiated in late mid-life improved measures of spatial memory in male mice. GeroScience 45, 2481–2494 (2023).

32. Stubbs, B. J. et al. In vitro stability and in vivo pharmacokinetics of the novel ketogenic ester, bis hexanoyl (R)-1,3-butanediol. Food and Chemical Toxicology 147, 111859 (2021).

33. Lopez-Rodriguez, A. B. et al. Acute systemic inflammation exacerbates neuroinflammation in Alzheimer’s disease: IL-1β drives amplified responses in primed astrocytes and neuronal network dysfunction. Alzheimer’s & Dementia 17, 1735–1755 (2021).

34. Weir, J. B. D. B. New methods for calculating metabolic rate with special reference to protein metabolism. J Physiol 109, 1–9 (1949).

35. Banks, W. A. et al. Lipopolysaccharide-induced blood-brain barrier disruption: roles of cyclooxygenase, oxidative stress, neuroinflammation, and elements of the neurovascular unit. J Neuroinflammation 12, 223 (2015).

36. Sumbria, R. K. et al. Aging exacerbates development of cerebral microbleeds in a mouse model. J Neuroinflammation 15, 69 (2018).

37. Lau, W. L. et al. Chronic Kidney Disease Increases Cerebral Microbleeds in Mouse and Man. Transl. Stroke Res. 11, 122–134 (2020).

38. Yoo, J.-Y., et al. LPS-Induced Acute Kidney Injury Is Mediated by Nox4-SH3YL1. Cell Reports 33, (2020).

39. Pinho, T. S., Verde, D. M., Correia, S. C., Cardoso, S. M. & Moreira, P. I. *O*-GlcNAcylation and neuronal energy status: Implications for Alzheimer’s disease. Ageing Research Reviews 46, 32–41 (2018).

40. Neudorf, H., Myette-Côté, É. & P. Little, J. The Impact of Acute Ingestion of a Ketone Monoester Drink on LPS-Stimulated NLRP3 Activation in Humans with Obesity. Nutrients 12, 854 (2020).

41. Agwunobi, A. O., Reid, C., Maycock, P., Little, R. A. & Carlson, G. L. Insulin resistance and substrate utilization in human endotoxemia. J Clin Endocrinol Metab 85, 3770–3778 (2000).

42. Wang, A. et al. Opposing Effects of Fasting Metabolism on Tissue Tolerance in Bacterial and Viral Inflammation. Cell 166, 1512–1525.e12 (2016).

43. Sens, J. et al. Lipopolysaccharide administration induces sex-dependent behavioural and serotonergic neurochemical signatures in mice. Pharmacol Biochem Behav 153, 168–181 (2017).

44. Cai, K. C. et al. Age and sex differences in immune response following LPS treatment in mice. Brain Behav Immun 58, 327–337 (2016).

45. Gonzalez, P. S. et al. Mannose impairs tumour growth and enhances chemotherapy. Nature 563, 719–723 (2018).

46. Liang, R. et al. PMI-controlled mannose metabolism and glycosylation determines tissue tolerance and virus fitness. Nat Commun 15, 2144 (2024).

47. Clarke, D. D. & Sokoloff, L. Substrates of Cerebral Metabolism. in Basic Neurochemistry: Molecular, Cellular and Medical Aspects*. 6th edition* (Lippincott-Raven, 1999).

48. Sloviter, H. a. & Kamimoto, T. The isolated, perfused rat brain preparation metabolizes manmose but not maltose. Journal of Neurochemistry 17, 1109–1111 (1970).

49. Kizer, D. E. & Mccoy, T. A. Phosphomannose isomerase activity in a spectrum of normal and malignant rat tissues. Proc Soc Exp Biol Med 103, 772–774 (1960).

50. Rastedt, W., Blumrich, E.-M. & Dringen, R. Metabolism of Mannose in Cultured Primary Rat Neurons. Neurochem Res 42, 2282–2293 (2017).

51. Dringen, R., Bergbauer, K., Wiesinger, H. & Hamprecht, B. Utilization of mannose by astroglial cells. Neurochem Res 19, 23–30 (1994).

52. Page, K. A. et al. Effects of Fructose vs Glucose on Regional Cerebral Blood Flow in Brain Regions Involved With Appetite and Reward Pathways. JAMA 309, 63–70 (2013).

53. Cha, S. H., Wolfgang, M., Tokutake, Y., Chohnan, S. & Lane, M. D. Differential effects of central fructose and glucose on hypothalamic malonyl–CoA and food intake. Proceedings of the National Academy of Sciences 105, 16871–16875 (2008).

54. Lynch, C. J. & Adams, S. H. Branched-chain amino acids in metabolic signalling and insulin resistance. Nat Rev Endocrinol 10, 723–736 (2014).

55. Xiao, M., Liu, D., Xu, Y., Mao, W. & Li, W. Role of PFKFB3-driven glycolysis in sepsis. Annals of Medicine 55, 1278–1289 (2023).

56. Trefely, S. et al. Kinome Screen Identifies PFKFB3 and Glucose Metabolism as Important Regulators of the Insulin/Insulin-like Growth Factor (IGF)-1 Signaling Pathway. J Biol Chem 290, 25834–25846 (2015).

57. Jimenez-Blasco, D. et al. Weak neuronal glycolysis sustains cognition and organismal fitness. Nat Metab 6, 1253–1267 (2024).

58. Yu, S. B. et al. Neuronal activity-driven O-GlcNAcylation promotes mitochondrial plasticity. Developmental Cell 59, 2143–2157.e9 (2024).

59. Hardie, D. G. & Carling, D. The AMP-Activated Protein Kinase. European Journal of Biochemistry 246, 259–273 (1997).

60. Min, S. H., Song, D. K., Lee, C. H., Roh, E. & Kim, M.-S. Hypothalamic AMP-Activated Protein Kinase as a Whole-Body Energy Sensor and Regulator. Endocrinol Metab 39, 1–11 (2024).

61. Shu, H. et al. Acute Nicotine Treatment Alleviates LPS-Induced Impairment of Fear Memory Reconsolidation Through AMPK Activation and CRTC1 Upregulation in Hippocampus. Int J Neuropsychopharmacol 23, 687–699 (2020).

62. Santos, G. A. et al. Hypothalamic AMPK activation blocks lipopolysaccharide inhibition of glucose production in mice liver. Molecular and Cellular Endocrinology 381, 88–96 (2013).

63. Suissa, L. et al. Ingested Ketone Ester Leads to a Rapid Rise of Acetyl-CoA and Competes with Glucose Metabolism in the Brain of Non-Fasted Mice. Int J Mol Sci 22, 524 (2021).

64. Zhao, X. et al. The Relationship between Branched-Chain Amino Acid Related Metabolomic Signature and Insulin Resistance: A Systematic Review. Journal of Diabetes Research 2016, 2794591 (2016).

65. Semmler, A. et al. Sepsis causes neuroinflammation and concomitant decrease of cerebral metabolism. J Neuroinflammation 5, 38 (2008).

66. Bellaver, B. et al. Activated peripheral blood mononuclear cell mediators trigger astrocyte reactivity. Brain, Behavior, and Immunity 80, 879–888 (2019).

67. Hart, B. L. Biological basis of the behavior of sick animals. Neurosci Biobehav Rev 12, 123–137 (1988).

68. Harrison, N. A. et al. A Neurocomputational Account of How Inflammation Enhances Sensitivity to Punishments Versus Rewards. Biol Psychiatry 80, 73–81 (2016).

69. López-Gambero, A. J., Martínez, F., Salazar, K., Cifuentes, M. & Nualart, F. Brain Glucose-Sensing Mechanism and Energy Homeostasis. Mol Neurobiol 56, 769–796 (2019).

70. Yang, H., Shan, W., Zhu, F., Wu, J. & Wang, Q. Ketone Bodies in Neurological Diseases: Focus on Neuroprotection and Underlying Mechanisms. Front Neurol 10, 585 (2019).

71. Jensen, N. J., Wodschow, H. Z., Nilsson, M. & Rungby, J. Effects of Ketone Bodies on Brain Metabolism and Function in Neurodegenerative Diseases. International Journal of Molecular Sciences 21, 8767 (2020).

72. Bohnen, J. L. B., Albin, R. L. & Bohnen, N. I. Ketogenic interventions in mild cognitive impairment, Alzheimer’s disease, and Parkinson’s disease: A systematic review and critical appraisal. Front Neurol 14, 1123290 (2023).

73. Cunnane, S. et al. Brain fuel metabolism, aging, and Alzheimer’s disease. Nutrition 27, 3–20 (2011).

74. Kellar, D. & Craft, S. Brain insulin resistance in Alzheimer’s disease and related disorders: mechanisms and therapeutic approaches. The Lancet Neurology 19, 758–766 (2020).

75. Nitchingham, A. et al. Long-acting intranasal insulin for the treatment of delirium—a randomised clinical trial. Age Ageing 54, afaf276 (2025).

76. Matsuura, T. R., Puchalska, P., Crawford, P. A. & Kelly, D. P. Ketones and the Heart: Metabolic Principles and Therapeutic Implications. Circ Res 132, 882–898 (2023).

77. Castellano, C.-A. et al. Lower brain 18F-fluorodeoxyglucose uptake but normal 11C-acetoacetate metabolism in mild Alzheimer’s disease dementia. J Alzheimers Dis 43, 1343–1353 (2015).

78. Eap, B. et al. Ketone body metabolism declines with age in mice in a sex-dependent manner. 2022.10.05.511032 Preprint at 10.1101/2022.10.05.511032 (2022).

