## Supplemental data for "Ketone bodies mitigate against systemic inflammation-induced changes in brain energy metabolism and delirium-like deficits in aged mice"

Pierre-Louis Hollier et al.

**Supplementary material**

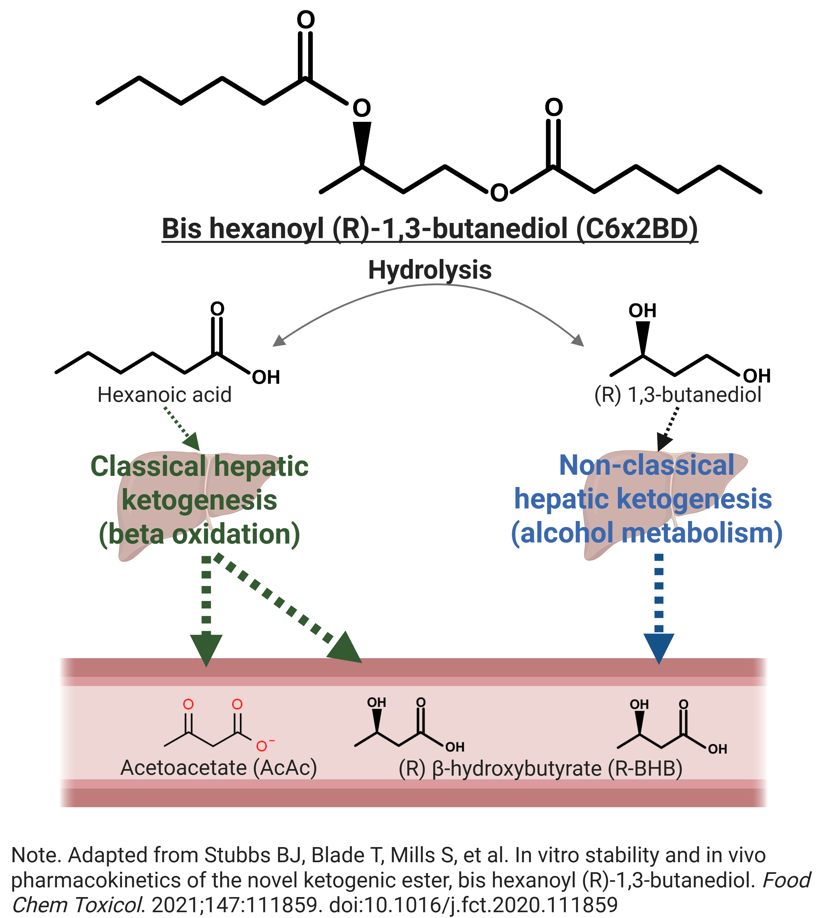

**Figure S1 Structure and metabolism of the synthetic ketone ester used in all experiments.**

C6x2BD is administered by oral gavage, hydrolysed to hexanoic acid or (R) 1,3-butanediol, both of which can be converted to β-hydroxybutyrate or acetoacetate in the liver and released into the blood.

**Table S1 List of primers for RT-PCR**

| **Gene** | **Forward** | **Reverse** | **Probe** |
| --- | --- | --- | --- |
| ***Aldob*** | AAGTGGCGTGCTGTGTTGAG | ATAGGGACCAGCCCATTCTGC | n/a |
| ***Bdh1*** | AGGCTGTGACTCTGGATTTGGG | CTGGATGGTTCTCAGTCGGTCA | n/a |
| ***Casp1*** | GGACAAGGCACGGGACCTA | GCTGATGGAGCTGATTGAAGCT | n/a |
| ***Cs*** | GTAGCTCTCTCCCTTCGGTC | ACGAGGCAGGATGAGTTCTTG | n/a |
| ***Cyc1*** | CAGCTTCCATTGCGGACAC | GGCACTCACGGCAGAATGAA | n/a |
| ***Eno1*** | AGTACGGGAAGGACGCCACCA | GCGGCCACATCCATGCCGAT | n/a |
| ***Gapdh*** | TGACCTCAACTACATGGTCTACA | CTTCCCATTCTCGGCCTTG | n/a |
| ***Gpi1*** | AGGAGACCATCACCAATGCAG | TCTTTCACTTTGGCCGTGTTC | n/a |
| ***Hk1*** | CCGCCATTGAAACGGATAAGG | TTGGCTGATCGGAAGGAGAC | n/a |
| ***Hk2*** | ACCAAGTGCAGAAGGTTGAC | TGCTGTAGGGTGTGTGGTAG | n/a |
| ***Il1b*** | GCACACCCACCCTGCA | ACCGCTTTTCCATCTTCTTCTT | TGGAGAGTCTGGATCCCAAGCAATACCC |
| ***Khk*** | AGGCAACGCATCCAACTC | CCAGGACAAAATCGGCAAC | n/a |
| ***Ldha*** | ACGCAGACAAGGAGCAGTGGAA | ATGCTCTCAGCCAAGTCTGCCA | n/a |
| ***Ldhb*** | CCTCAGATCGTCAAGTACAGCC | ATCCGCTTCCAATCACACGGTG | n/a |
| ***Mpc1*** | CTCCAGAGATTATCAGTGGGCG | CTCCAGAGATTATCAGTGGGCG | n/a |
| ***Mpc2*** | ACCACCGACTCATGGATAAAGTG | AGCACACACCAATCCCCATTTC | n/a |
| ***Mpi*** | CACACCCTAACAAGGAGCTGG | AGCCCTGGAAGGAGGTAAGG | n/a |
| ***Nfe2l2*** | AGCAACTCCAGAAGGAACAGG | AATGTGGGCAACCTGGGAG | n/a |
| ***Nlrp3*** | GAGCCTACAGTTGGGTGAAATGT | CCACGCCTACCAGGAAATCTC | n/a |
| ***Oxct1*** | GAGCGACAGTTCCTTTCTGGTG | TACACCAGCACAGGGTATGGGA | n/a |
| ***Pdk1*** | TCCCCCGATTCAGGTTCAC | CCCGGTCACTCATCTTCACA | n/a |
| ***Pfkfb3*** | TGTCCTGAAACTGACACCTG | AGCTCTTCATGTTCTCTGACC | n/a |
| ***Pfkp*** | CAGTCAGTGCCAACATAACCAA | CGGGATGCAGAGCTCATCA | n/a |
| ***Pgam1*** | ATGAGCGACACTATGGCGG | TGCGTACCTGCGATCCTTG | n/a |
| ***Pgk1*** | GGAGGCCCGGCATTCTGCAC | AGTCCACCCTCATCACGACCCG | n/a |
| ***Prkaa2*** | CAGAAGATTCGCAGTTTAGATGTTG | ACCTCCAGACACATATTCCATTACC | n/a |
| ***Slc16a1*** | CATTGGTGTTATTGGAGGTC | GAAAGCCTGATTAAGTGGAG | n/a |
| ***Slc16a3*** | TCAATCATGGTGCTGGGACT | TGTCAGGTCAGTGAAGCCAT | n/a |
| ***Slc16a7*** | CACCACCTCCAGTCAGARCG | CTCCCACTATCACCACAGGC | n/a |
| ***Slc2a1*** | CATCAACAGAGGGTGCCAACA | CACTGCTCCCAGGATGACAC | n/a |
| ***Slc2a2*** | ATCAACATGATCTTCACGGCTGT | TGGCAGTCATGCTCACGTAACTC | n/a |
| ***Tnf*** | CTCCAGGCGGTGCCTATG | GGGCCATAGAACTGATGAGAGG | TCAGCCTCTTCTCATTCCTGCTTGTGG |

**Figure S2**

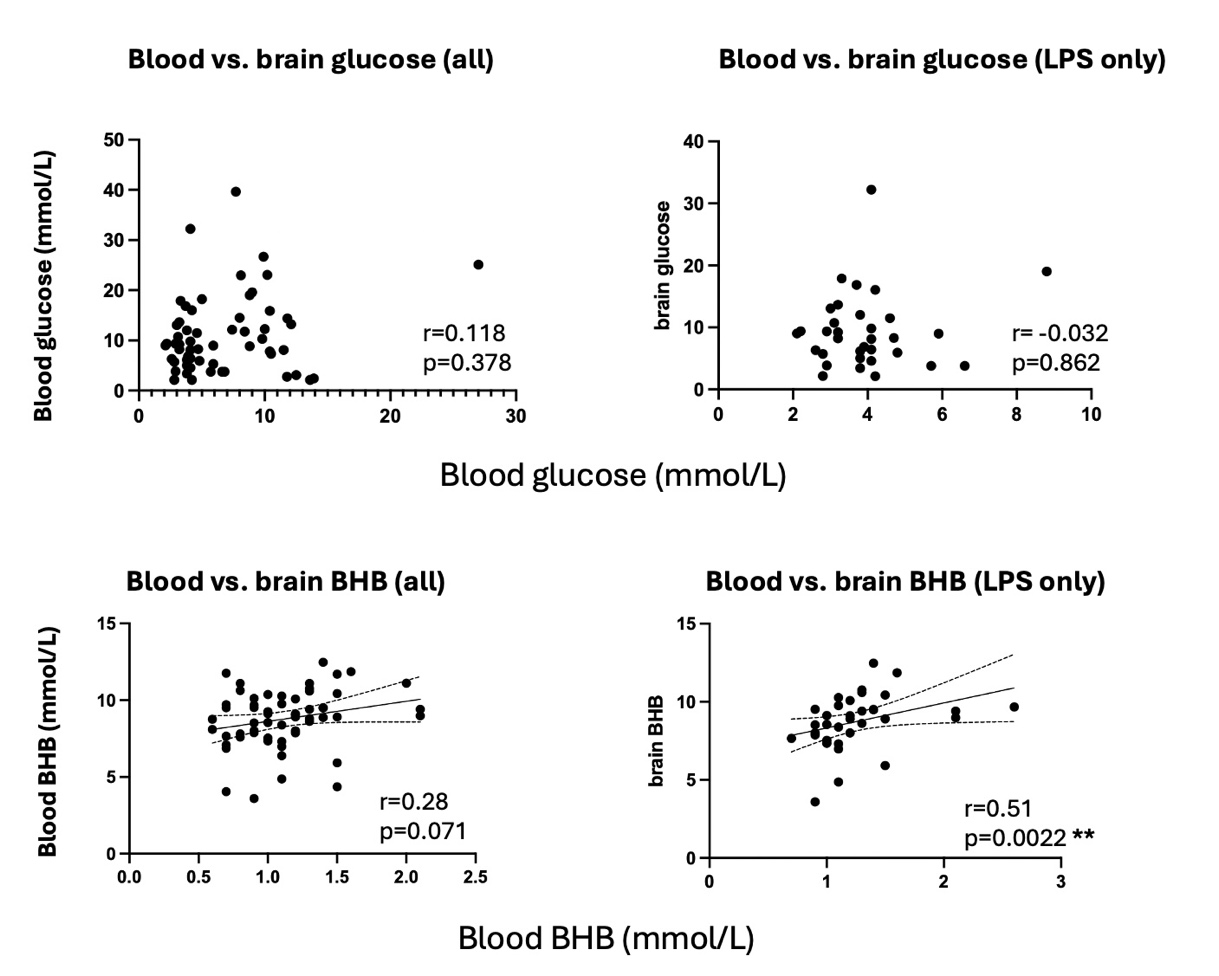

**Figure S2 Relationship of blood metabolite levels to brain metabolite levels.**

Correlations were performed using all animals treated with LPS (250 µg/Kg) or saline and with or without KE (3g/Kg) for which both blood and brain levels of glucose and BHB were measured. In the graphs on the left these samples were all included (n=58) while on the right only samples from animals treated with LPS were included (n=33). Linear Regression and resulting Pearson’s r are shown. The data show that blood glucose does not significantly influence brain glucose in either the full data set nor in the LPS only data. Conversely Brain BHB is influenced by blood BHB levels, particularly in the case of the LPS-treated animals.

**Figure S3. Metabolic enzymes affected by LPS and/or KE**

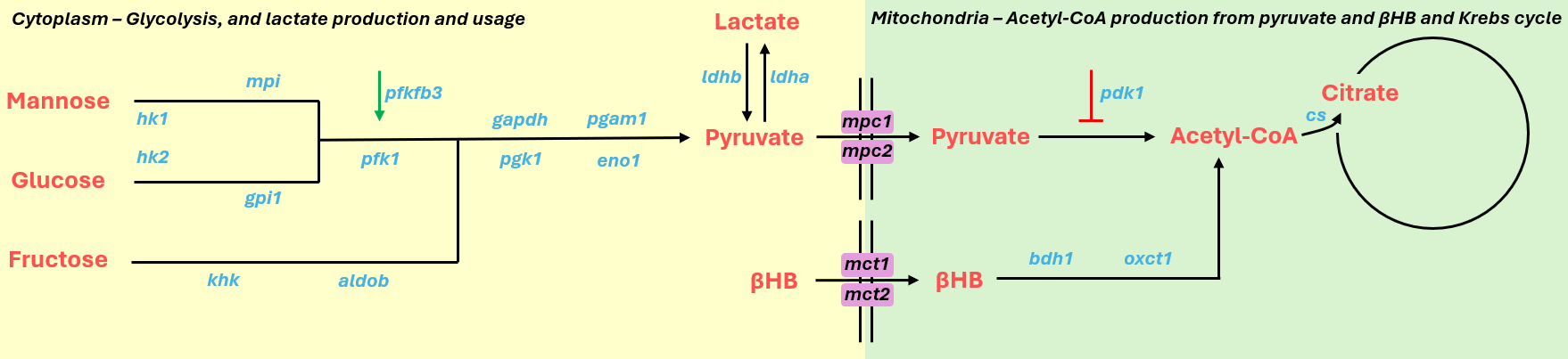

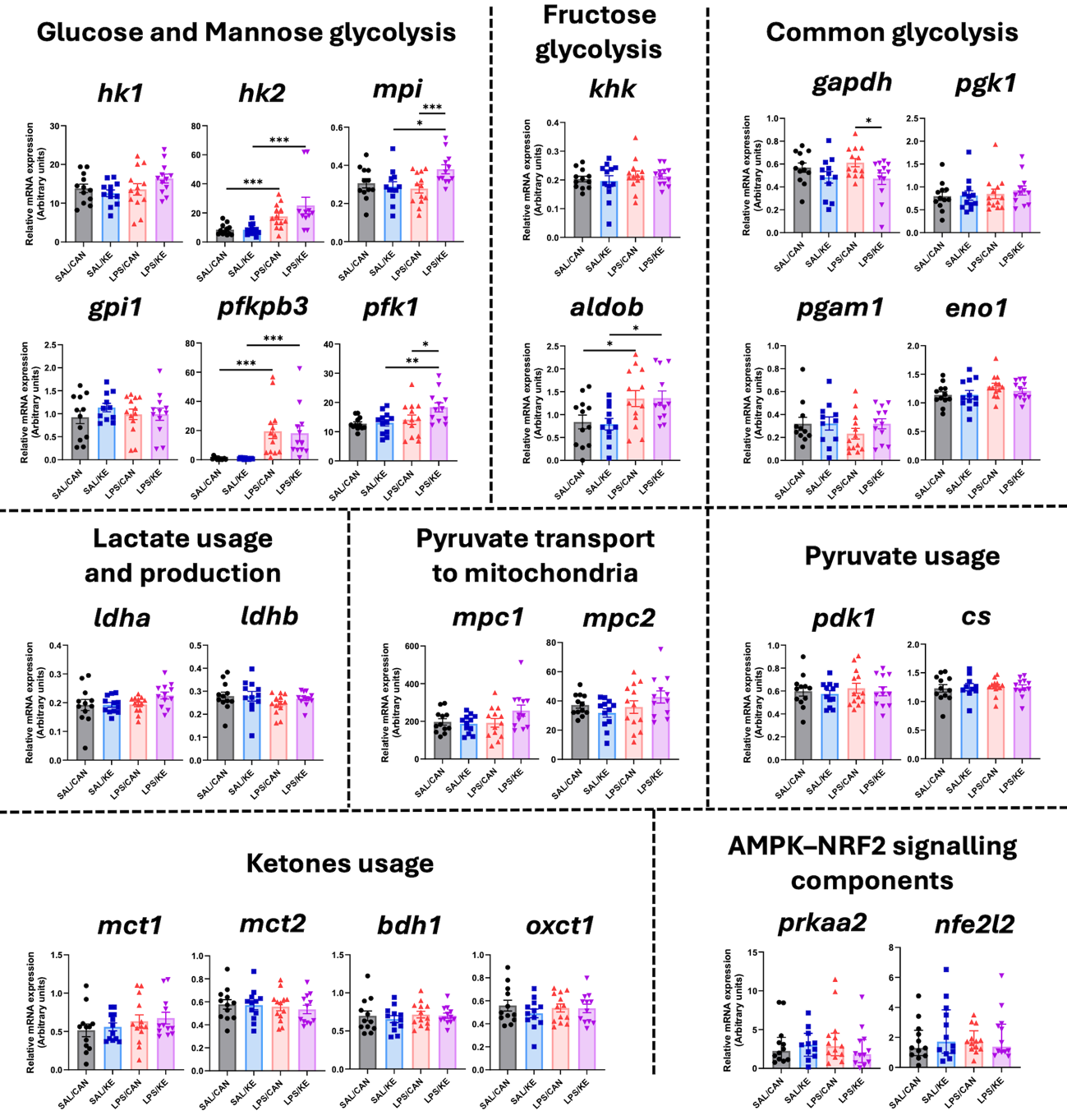

**Supplementary Figure 3. Effect of LPS and KE interventions on metabolic enzymes in hippocampus.** Graphs show transcripts of enzymes and transporters in these processes in the hippocampi of mice treated with LPS (250 μg/kg) or SAL as control and intervened with KE (3g/kg) or CAN as control, n=11-13/group. Fold increases in the transcripts were compared using 2-way ANOVA with treatment and intervention as between subjects factors. Statistically significant differences by post-hoc pairwise comparisons are denoted by ∗ (p < 0.05), ∗∗ (p < 0.01), and ∗∗∗ (p < 0.001). Normal data points are displayed as mean ± SEM whereas nonparametric data points are displayed as median ± IQ.

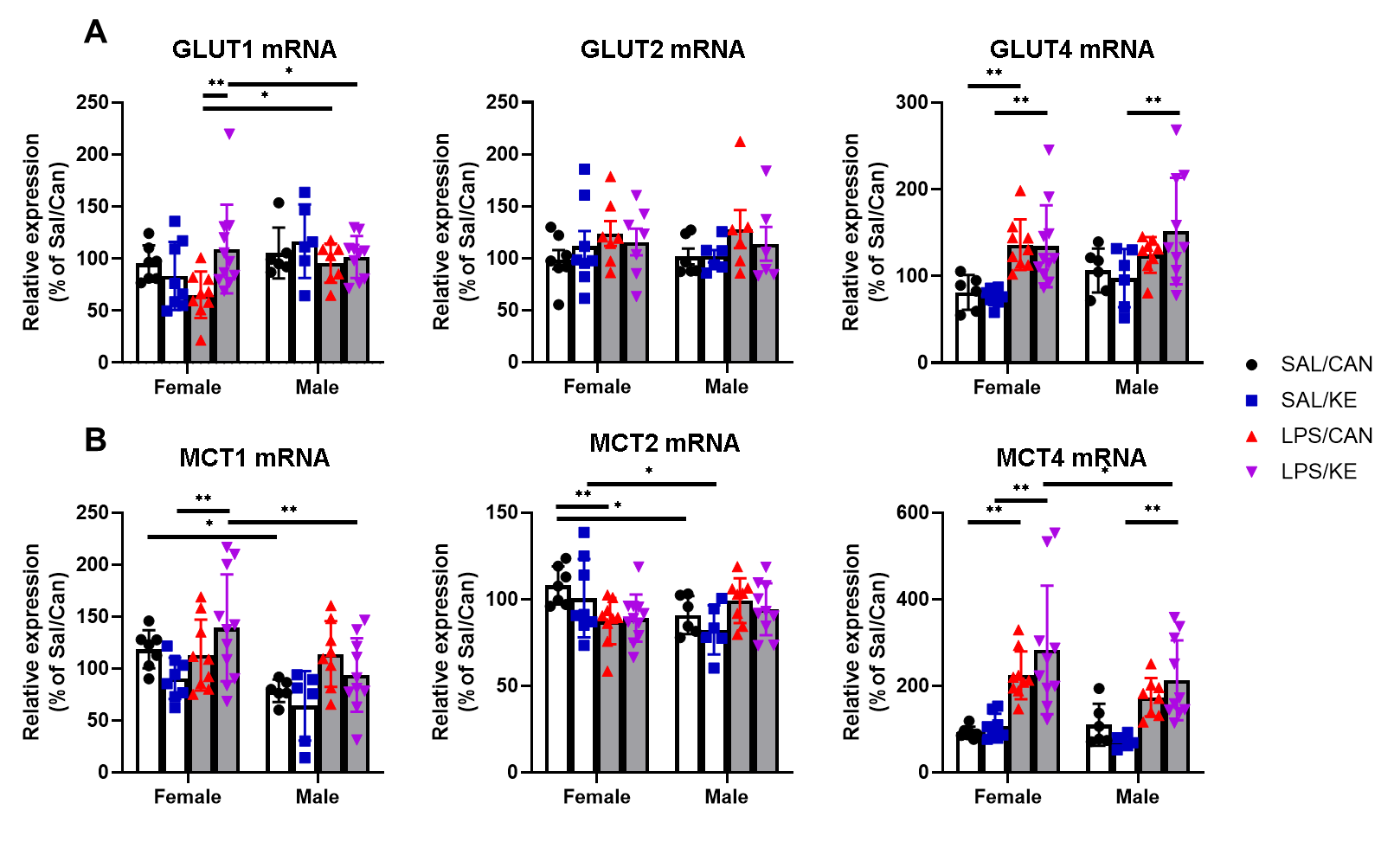

**Figure S4. mRNA expression of glucose and monocarboxylate transporters.**

Aged mice (20-25 months) were injected with 250µg/kg LPS i.p. or 10mL/kg saline, followed at 1 hour by KE or canola (3g/Kg by oral gavage). Hippocampal expression of transcripts for glucose and monocarboxylate transporters were assessed by RT-qPCR. LPS significantly increased the expression of Glut4 (F_1,56_=23.66, P<0.0001), MCT4 (F_1,56_=42.79, P<0.0001), and MCT1 (F_1,57_=9.920, P=0.0026) mRNAs. MCT1 expression was also increased in female (F_1,57_=10,.42 P<0.0021). Glut1 was on the other hand increased in males (F_1,56_=5.04, P=0.0287) and an interaction of sex, LPS and KE treatment was found (F_1,56_=4.295, P=0.0428). SAL/CAN n= 7♀/6♂; SAL/KE 8♀/6♂; LPS/CAN 9♀/8♂; LPS/KE 11♀/10♂.

SAL: Saline; CAN: Canola oil; LPS: Lipopolysaccharide; KE: Ketone ester. Data are shown as mean ±SEM.
